# Multifaceted Representation of Genes via Deep Learning of Gene Expression Networks

**DOI:** 10.1101/2024.03.07.583777

**Authors:** Zheng Su, Mingyan Fang, Andrei Smolnikov, Marcel E. Dinger, Emily C. Oates, Fatemeh Vafaee

**Affiliations:** School of Biotechnology and Biomolecular Sciences, Faculty of Science, The University of New South Wales, Sydney, NSW 2052, Australia; BGI Research, Shenzhen 518083, China; BGI Australia, Herston, Queensland, Australia; School of Life and Environmental Sciences, Faculty of Science, University of Sydney, Sydney, NSW 2006, Australia

**Author notes:** These authors contributed equally to this work (Zheng Su, Mingyan Fang). These corresponding authors contributed equally to this work.

## Abstract

Accurate predictive modeling of human gene relationships would fundamentally transform our ability to uncover the molecular mechanisms that underpin key biological and disease processes. Recent studies have employed advanced AI techniques to model the complexities of gene networks using large gene expression datasets^1–11^. However, the extent and nature of the biological information these models can learn is not fully understood. Furthermore, the potential for improving model performance by using alternative data types, model architectures, and methodologies remains underexplored. Here, we developed GeneRAIN models by training on a large dataset of 410K human bulk RNA-seq samples, rather than single-cell RNA-seq datasets used by most previous studies. We showed that although the models were trained only on gene expression data, they learned a wide range of biological information well beyond gene expression. We introduced GeneRAIN-vec, a state-of-the-art, multifaceted vectorized representation of genes. Further, we demonstrated the capabilities and broad applicability of this approach by making 4,797 biological attribute predictions for each of 13,030 long non-coding RNAs (62.5 million predictions in total). These achievements stem from various methodological innovations, including experimenting with multiple model architectures and a new ‘Binning-By-Gene’ normalization method. Comprehensive evaluation of our models clearly demonstrated that they significantly outperformed current state-of-the-art models^3,12^. This study improves our understanding of the capabilities of Transformer and self-supervised deep learning when applied to extensive expression data. Our methodological advancements offer crucial insights into refining these techniques. These innovations are set to significantly advance our understanding and exploration of biology.

## Introduction

Gene function and its correlation with phenotypes is complex, and is encoded by a network of interdependent interactions and regulatory processes. Understanding and modeling these intricate relationships is key to addressing numerous biological questions and advancing biomedical applications including new treatment development. The advent of deep learning, recently accelerated by the introduction of Transformer models^13^ and the application of self-supervised training^14–16^, has had a significant impact on a broad spectrum of fields, including research, industry, and various aspects of daily life^17–20^. Attempts to harness these advanced technologies to better understand gene relationships through gene expression data have shown great promise^1,2^. These approaches have been used successfully to advance cell type classification, disease gene prediction, and epigenetic modification prediction^3–6^. These models have also been employed to simulate transcriptomic responses to perturbations and drug target identification^7–11^.

However, our understanding of the extent and nature of the biological information learned by these models is limited. The potential applications of this information have also not been fully explored. Furthermore, to date, most models have been trained on single-cell RNA-seq data, and the use of bulk RNA-seq data for such models and the performance of such models remains largely unexplored. In addition, while Bidirectional Encoder Representations from Transformers (BERT) models^15^, which reconstruct information of randomly masked genes using other genes, have been widely used, the effectiveness of other architectures like the Generative Pre-Training (GPT) model^16^, known for its success in large language models^21^, has not been thoroughly investigated.

In this study, we aim to address these gaps by comprehensively evaluating the capabilities of these models in learning diverse biological attributes of genes and simulating transcriptomic responses to genetic perturbations. We also explore strategies to improve model performance and investigate potential applications of the information learned by the models. Our goal is to deepen our understanding of the characteristics of these models and the nature of the information they learn, contributing to methodological innovations and the development of new applications.

## Results

### Self-supervised deep learning on transcriptomes

To explore the learning capabilities of deep learning models on large gene expression data, we began by training a model using a dataset of 722,425 human bulk RNA-seq samples from the ARCHS4 database^22^. After filtering for high-quality samples and removing potential single-cell samples (Methods), 410,850 samples remained for training our model. These samples represent a broad spectrum of phenotypes, tissues, cell lines, ages, and genders. We aimed to extract biological information from the constraints and interrelationships in expression among genes of each individual transcriptome, in the absence of phenotype data. We achieved this by randomly selecting 15% of genes in each sample and had their identities (gene symbols) masked. A six-layer BERT model^15^ (Fig. 1a) was then trained to reconstruct the identities of the masked genes by learning the interrelationship between genes (Extended Data Fig. 2a). The training data was prepared by computing gene expression z-scores for each gene (Methods and Extended Data Fig. 1a). Genes in each sample were subsequently sorted from highest to lowest based on their expression z-scores. In each sample, gene expression, represented by the gene order, was vectorized and added to gene identity vectors.

**Fig. 1.**
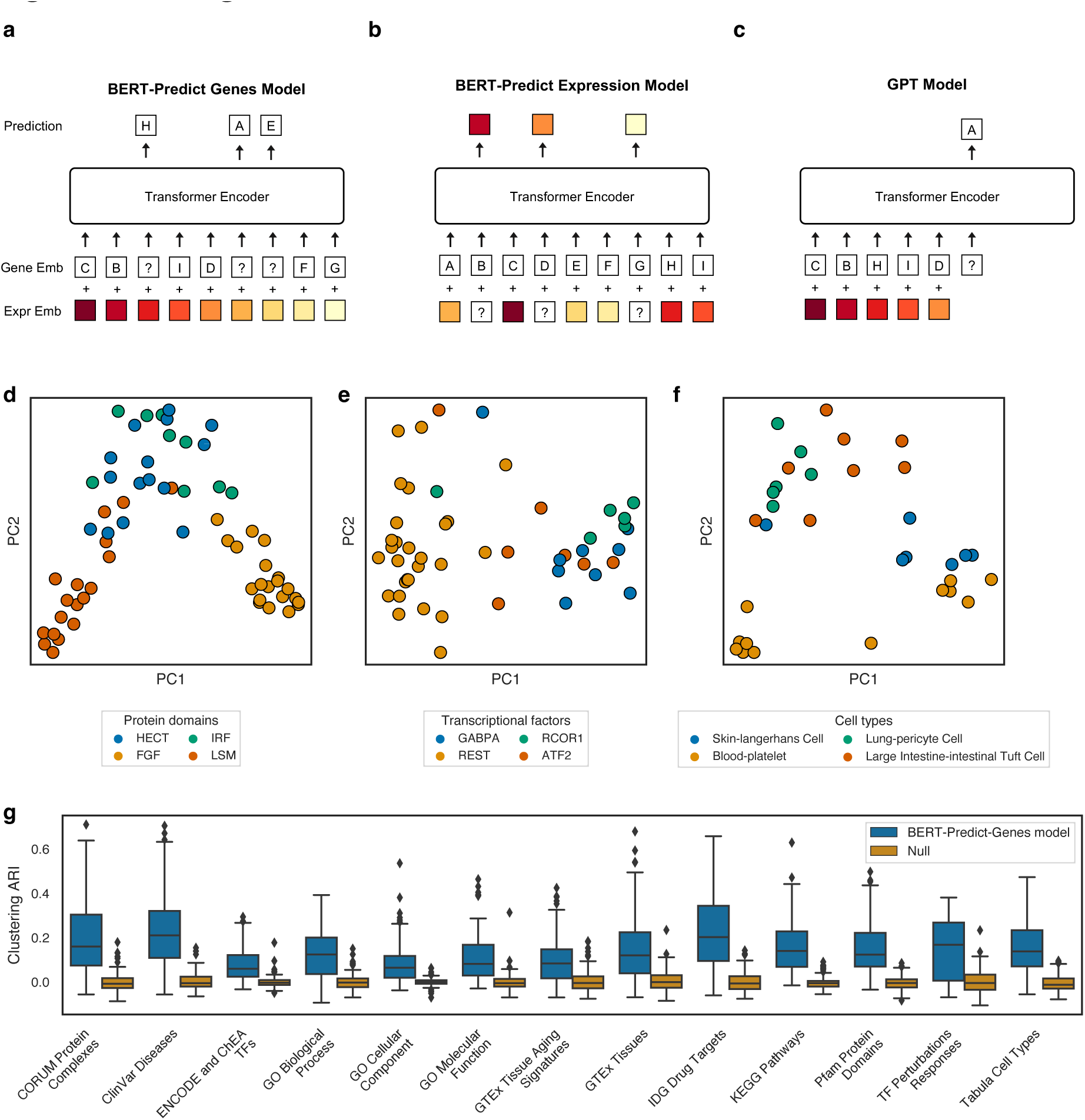
| Overview of model architectures and gene attributes learned by the BERT-Predict-Genes model. **a**, BERT Predict Genes model: Gene and expression embeddings are represented by bottom two rows of squares, with gene identities denoted by letters and expression levels by color intensity. Genes are sorted by expression in descending order; identities (gene symbol) of randomly selected genes are masked, and are predicted by Transformer encoder. **b,** BERT Predict Expression model: The model is trained to predict masked expressions rather than gene identities. **c,** GPT model: Genes are ordered by expression for next-gene prediction based on prior higher-expressed genes’ identities. **d-f,** PCA visualizations of gene embeddings from the BERT-Predict-Genes model, illustrating attributes of the protein domains of encoding proteins from the Pfam database, targeted transcription factors from ENCODE database and cell type markers from Tabula Sapiens database. **g,** Boxplot demonstrating the clustering agreement of gene embeddings from our BERT-Predict-Genes model with biological attributes of genes across various databases, assessed via the ARI against a null permutation distribution. Each box contains 100 data points, each from an experiment of random selections of four gene groups from the database. Center line, median; box limits, upper and lower quartiles; whiskers, 1.5× interquartile range; outliers, points. In all comparisons, p-values were < 1E-10, determined by a two-sided Mann-Whitney U test with Benjamini-Hochberg adjustment.

Following training, we investigated the information learned by the model. Gene embeddings, which are vectors of 200 numbers, represent the roles of each gene learned from the gene expression relationship. They capture a significant amount of information, as more than half (51.4%, Supplementary Table 1) of the model’s parameters resided within them. Therefore, we assessed how well the gene embeddings could represent a variety of biological attributes of genes, to understand what biological information the model had learned. For example, we explored whether gene embeddings contained information about the domains of proteins encoded by genes. This was tested by randomly selecting four protein domains from the Pfam database^23^, each of which was encoded by a specific group of genes. We then analyzed if embeddings of these genes in our model could recapitulate these groupings. The amount of information learned by the model was measured by the performance of the recapitulation, accessed via clustering Adjusted Rand Index (ARI)^24,25^ (Methods). The analysis revealed that gene embeddings had captured protein domain information, as illustrated in Fig. 1d, g.

In addition to protein domains, we evaluated multiple other biological attributes of genes. These included protein-protein interactions from the CORUM database, gene-disease associations from the ClinVar database, transcription factor targets from the ENCODE database and ChIP-seq Enrichment Analysis (ChEA) dataset, Gene Ontology (GO) attributes (including biological processes, cellular components, and molecular functions)^26^, tissue aging signatures from the GTEx database, GTEx tissue associations, drug targets from the IDG database, pathways from the KEGG database, gene expression responses to transcription factor (TF) perturbations (Fig. 1e), and cell type-specific markers from the Tabula database (Methods and Fig. 1f). For each of these attributes we found that the gene embeddings in our model exhibited learning capabilities (Fig. 1g, Extended Data Fig. 3 and Supplementary Table 2). This suggests that although the model was trained on only expression data, it could learn biological information beyond expression.

### Normalization methods

During our analysis, we observed potential bias being introduced by the initial data normalization method. We noted that the highest-ranking genes in a sample often had extremely high z-scores. This was primarily due to their low mean expression across samples, resulting in them appearing as outliers in the gene expression distribution (e.g., gene A and C in Extended Data Fig.1a). Conversely, genes with expression patterns closer to normal distributions rarely ranked highly in a sample (e.g., gene E in Extended Data Fig. 1a). This bias could risk confounding the gene expression rankings in a sample, skewing high ranking genes toward those with atypical expression distributions, while genes with normal distributions were less likely to be top-ranked. This could potentially degrade the model’s performance as only the top-ranked genes would be selected and inputted into the model. To mitigate this issue, we employed a novel normalization strategy, termed the ‘Binning-By-Gene’ method. In this method, for each gene, expressions across samples were allocated into one of 2,000 bins based on their expression rank. Genes with no expression value were allocated to the lowest bin, while non-zero expressions were evenly distributed across other bins (Methods, Extended Data Fig. 1b). This normalization method equalized the probability of each gene occupying any rank position in the model input, thus reducing the bias we observed in the Z-Score-based method.

### Gene Attribute Learning Index

To facilitate a comprehensive evaluation of the performance of various models, and models utilizing different normalization methods, in learning gene biological attributes within their embeddings, we developed the Attribute Learning Index. This index is an average of clustering consistency metrics; ARI, Fowlkes-Mallows Index (FMI)^27^, and Normalized Mutual Information (NMI)^28^, calculated between the model embedding-based clustering and the actual gene biological attribute groupings, compared to random. We used three different metrics to reduce the bias from a single metric. The index averages values from 100 random selections of four groups for clustering comparisons. To minimize potential biases from gene attribute databases, we further included databases such as Azimuth (cell type markers), BioPlex (protein complexes), HumanCyc (metabolite pathways), InterPro (Protein Domains), MSigDB Hallmark (biological states/processes), OMIM (disease association), huMAP (protein complex), and lncHUB (lncRNA co-expression) (detailed in Supplementary Table 3 and Methods). This approach provided a comprehensive evaluation of the model’s capability in learning biological attributes of genes. Using the Attribute Learning Index, we compared our normalization method, and found that our ‘Binning-By-Gene’ normalization method significantly enhanced the model’s efficiency in learning gene biological attributes (p = 0.007 by t-test, Fig. 2a and Supplementary Table 2, BERT-Pred-Genes model).

**Fig. 2.**
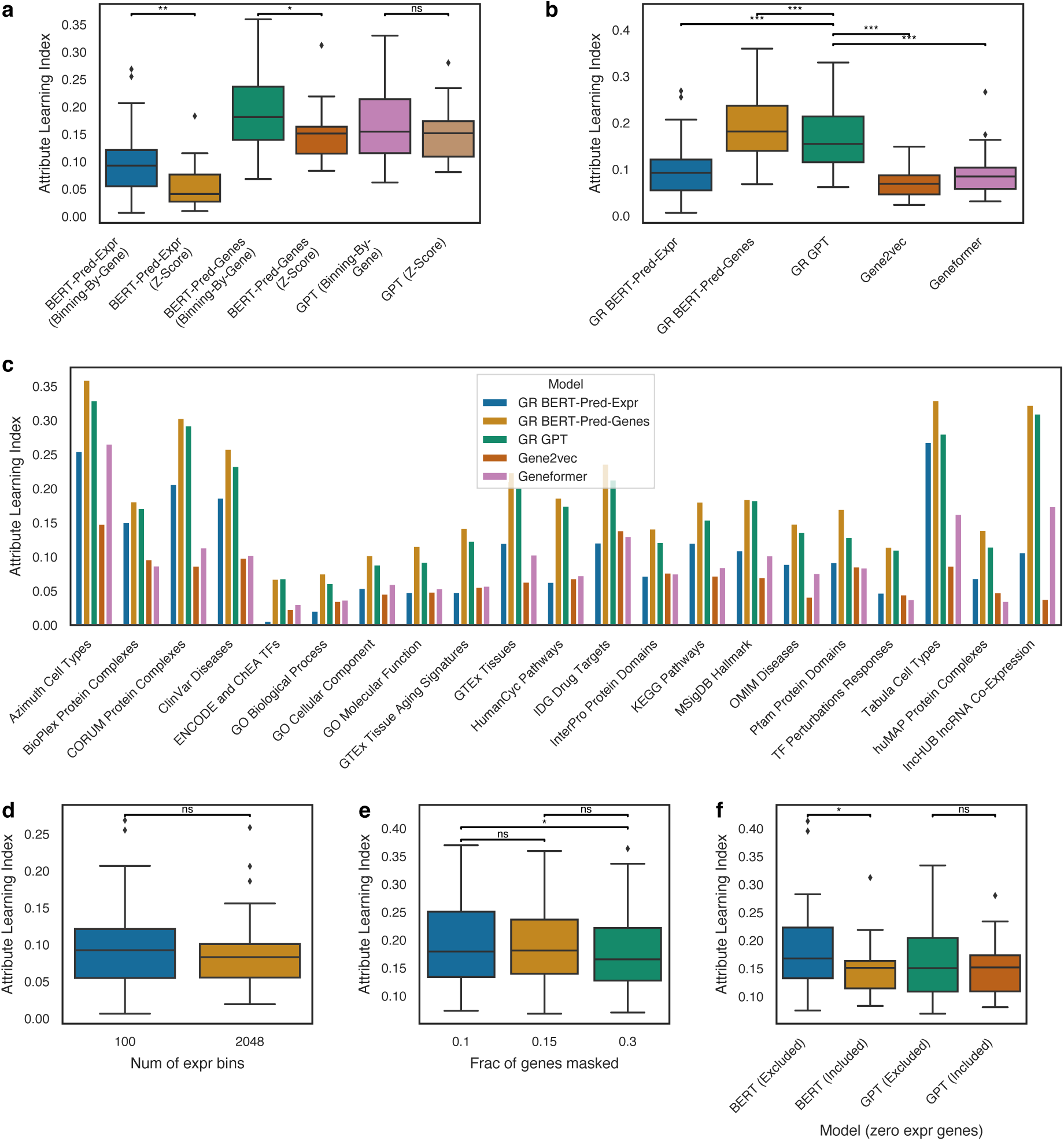
| Comparative analysis of gene attribute learning ability. **a**, Comparisons of gene Attribute Learning Indices between models employing ‘Z-Score’ and ‘Binning-By-Gene’ normalization methods. **b,** Assessment of gene Attribute Learning Indices from different GeneRAIN (GR) models, Gene2vec, and Geneformer. ‘Binning-By-Gene’ method was used for GR models. **c,** Detailed breakdown of gene attribute databases for results in panel b. **d,** Analysis of gene Attribute Learning Indices in BERT-Pred-Expr models with different expression binning numbers. **e,** Investigation of gene Attribute Learning Indices in BERT-Pred-Genes models having different fractions of masked genes. **f,** Comparison of gene Attribute Learning Indices in BERT-Pred-Genes and GPT models, considering the inclusion or exclusion of zero expression genes in z-score normalization. Each of the boxplots in these figures represents 21 data points, each from a gene attribute database. Center line, median; box limits, upper and lower quartiles; whiskers, 1.5× interquartile range; outliers, points. A two-sided paired t-test was used for statistical comparisons, after confirming the normality of the data with Shapiro-Wilk tests. *** *p* < 0.00001, ** *p* < 0.001, * *p* < 0.05, ‘ns’ indicates not significant (*p* >= 0.05).

### Comparison of different model architectures and configurations

As variations in model architecture and configuration can significantly impact performance, we sought to improve the training method to maximize the model performance. In addition to our initial BERT model that masked gene identities (termed ‘BERT-Pred-Genes’), we explored another BERT model (‘BERT-Pred-Expr’), which masked gene expression instead of gene identities, after expressions were discretized into 100 bins. The model was trained to predict these masked expressions (Fig. 1b and Extended Data Fig. 2c, d). Additionally, we experimented with a GPT model^16^, where genes were ordered by expression from the highest to the lowest before input into the model. The model was trained to predict the next gene in a sequence based on its higher expression predecessors (Fig. 1c and Extended Data Fig. 2e, f). We collectively termed our models as ‘GeneRAINs’ (abbreviated as GR), reflecting their focus on gene representation, using AI and network-oriented approaches.

We assessed the gene attribute learning capabilities of the gene embeddings from our models, compared them with those of the state-of-the-art Geneformer model^3^, a BERT model pre-trained on 30M single-cell transcriptomes, and the widely-used gene vectors Gene2vec^12^. Our findings indicated that both the GR BERT-Pred-Genes and GR GPT models significantly outperformed Geneformer and Gene2vec models (p < 0.00001 by t-test, Fig. 2b, c and Supplementary Table 2). We also compared different normalization methods, observing that the ‘Binning-By-Gene’ approach showed significantly superior performance over the Z-Score method in our two BERT based models, although differences were less pronounced in the GPT model (Fig. 2a, Extended Data Fig. 4a and Supplementary Table 2).

Further analyses focused on the impact of various hyperparameters. In the BERT-Pred-Expr model, we observed that increasing the number of expression bins from 100 to 2048 did not have significant impact on model performance (p = 0.06 by t-test, Fig. 2d). We also found that the proportion of masked genes in the BERT-Pred-Genes model appeared to have small effects on performance (Fig. 2e). Furthermore, we assessed the impact of including or excluding zero-expression genes in the z-score normalization for both BERT-Pred-Genes and GPT models. Excluding zero-expression genes from z-score calculations slightly improved the performance of the BERT-Pred-Genes model (Fig. 2f). Lastly, we compared the gene Attribute Learning Index across training epochs and noted that GPT models exhibited a faster learning rate compared to BERT models (Extended Data Fig. 4b-d).

### Diverse representation of gene attributes in gene embeddings

As different biological attributes can be correlated, a possible concern is that multiple gene attributes learned by the model actually represent only a singular attribute. For instance, consider three gene attributes: pathway association, disease association, and protein-protein interaction of encoded proteins. They might represent a single attribute rather than three different ones, as proteins from the same complex could be more likely involved in similar pathways or associated with the same diseases. To investigate this possibility, we examined whether different gene attributes contained the same information, by determining if they are associated with the same dimensions of the gene embedding. Specifically, we measured the extent to which each dimension of the gene embedding explains the variability of each gene attribute. This was accomplished using the normalized F-statistic in an Analysis of Variance (ANOVA) analysis in each clustering of gene Attribute Learning Index calculation, with the F-statistic values indicating how much variation each gene embedding dimension explains with respect to the differences between gene attribute clusters. Thus, the F-statistic serves as a measure of the association between each gene attribute and each gene embedding dimension (Methods).

Our findings indicate that while there is some overlap in the dimensions associated with different gene attributes, the dimensions associated with different gene attributes are very diverse and diverse dimensions are associated with a single gene attribute in our models (Fig. 3a and Extended Data Fig. 5-7). This finding is in contrast with the Geneformer embeddings, which showed much less diversity (Fig. 3b). The diversity in our models suggests that these gene attributes are mainly distinct, and that the models were primarily learning different gene properties. This result underscores the ability of our models to capture a broad spectrum of gene attributes, reflecting their complex biological roles and interactions.

**Fig. 3.**
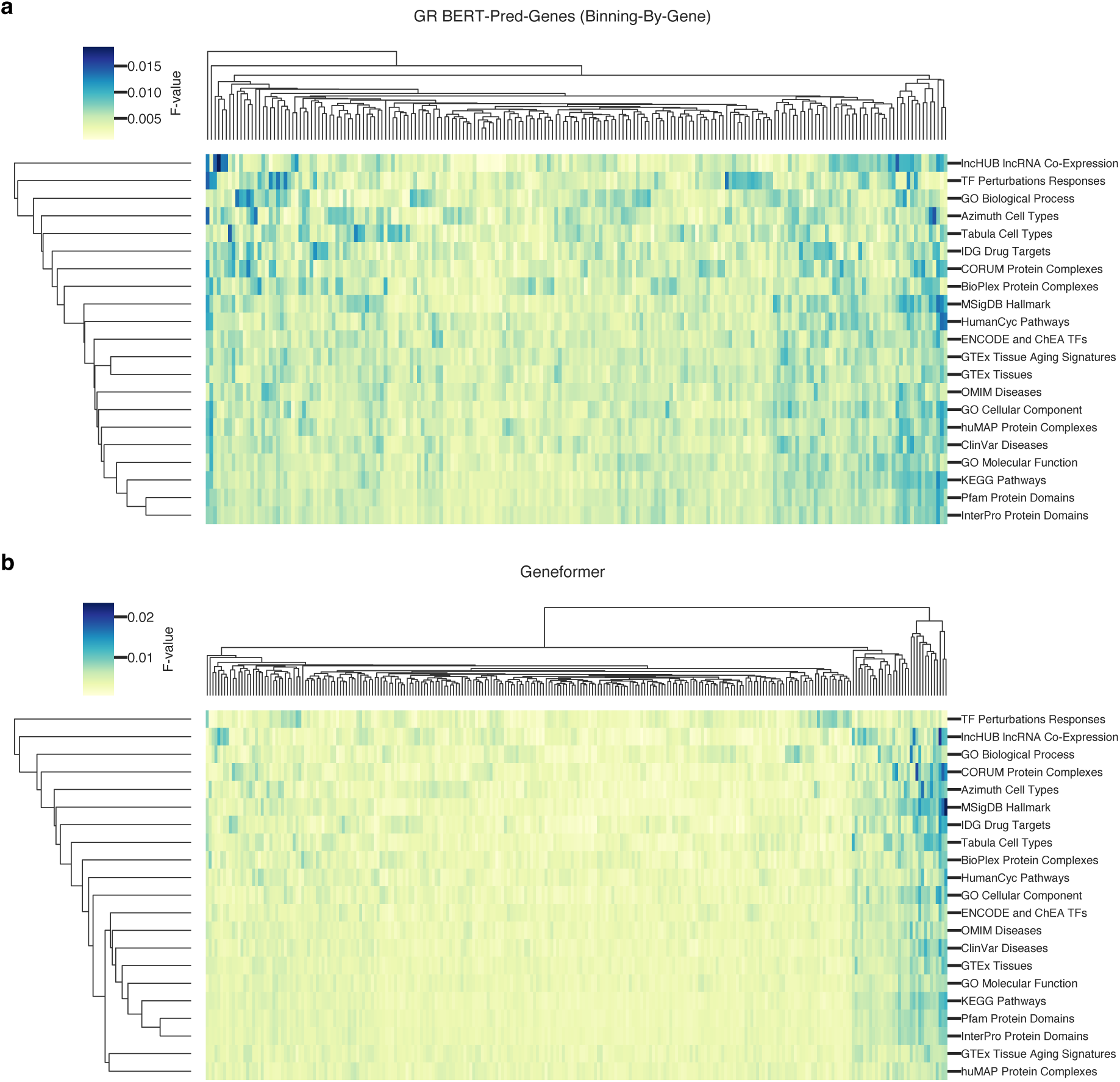
| Dimensional contributions in gene embeddings to gene attribute learning. This figure depicts the relative importance of each embedding dimension in clustering biological attributes of genes, as indicated by the color intensity representing the normalized F-statistic from ANOVA (Methods). **a,** Contribution distribution in the BERT-Pred-Genes model, with each row corresponding to gene attributes from a specific database and columns denoting embedding dimensions. **b,** The distribution for the Geneformer model.

### Models’ ability in learning transcriptomic responses to genetic perturbations

In addition to gene attribute learning, a robust model should also perform well in other tasks related to understanding gene relationships. Therefore, we assessed our models’ performance based on another important metric: their ability in learning about transcriptomic responses to genetic perturbations, such as the knockdown of specific genes. We investigated this by determining if genes that lead to similar perturbation transcriptome response also have similar encoding in the models. This aspect of model evaluation differs from our previous assessment of gene biological attributes, as it reflects the models’ ability in learning a gene’s regulatory influence on the expression of other genes. Successfully replicating transcriptomic responses requires the models to have learnt the intricate regulatory relationships among genes. This is likely to be also encoded within the Transformer encoder’s hidden layer parameters, rather than solely within the gene embedding layer.

To evaluate this aspect of our models’ performance, we utilized data from the Replogle et al. study^29^, one of the largest Perturbation-sequencing (Perturb-seq)^30,31^ experiments published to date. This study employed CRISPRi^32–34^ to generate genome-scale knockdowns of common essential genes, followed by single-cell RNA-sequencing to measure the resulting transcriptomic changes. The high specificity of CRISPRi technology greatly enhanced the quality of this dataset, providing a robust basis for our evaluation^35^.

For each gene in the Perturb-seq dataset, we identified the five other genes whose knockdown resulted in the most similar transcriptomic responses, using a k-Nearest Neighbor (kNN) algorithm. We then assessed each model’s ability to recapitulate these nearest neighbor relationships, by measuring the overlap in the neighbors from model’s encoding space with those from Perturb-seq experiment (Methods). The significance level of overlap was determined by comparing the results to null distributions from permutation of embeddings.

We found that all our models contained significant information about transcriptomic responses. Notably, gene embeddings from models employing the ‘Binning-By-Gene’ normalization method showed higher overlap in nearest neighbors compared to those using Z-Score normalization in both BERT-Pred-Genes and GPT models (Fig. 4a and Supplementary Table 4). Additionally, when comparing our models’ gene embeddings with those from Geneformer, all three of our models using ‘Binning-By-Gene’ normalization demonstrated greater performance in capturing the transcriptome response information (Fig. 4b and Supplementary Table 4).

**Fig. 4.**
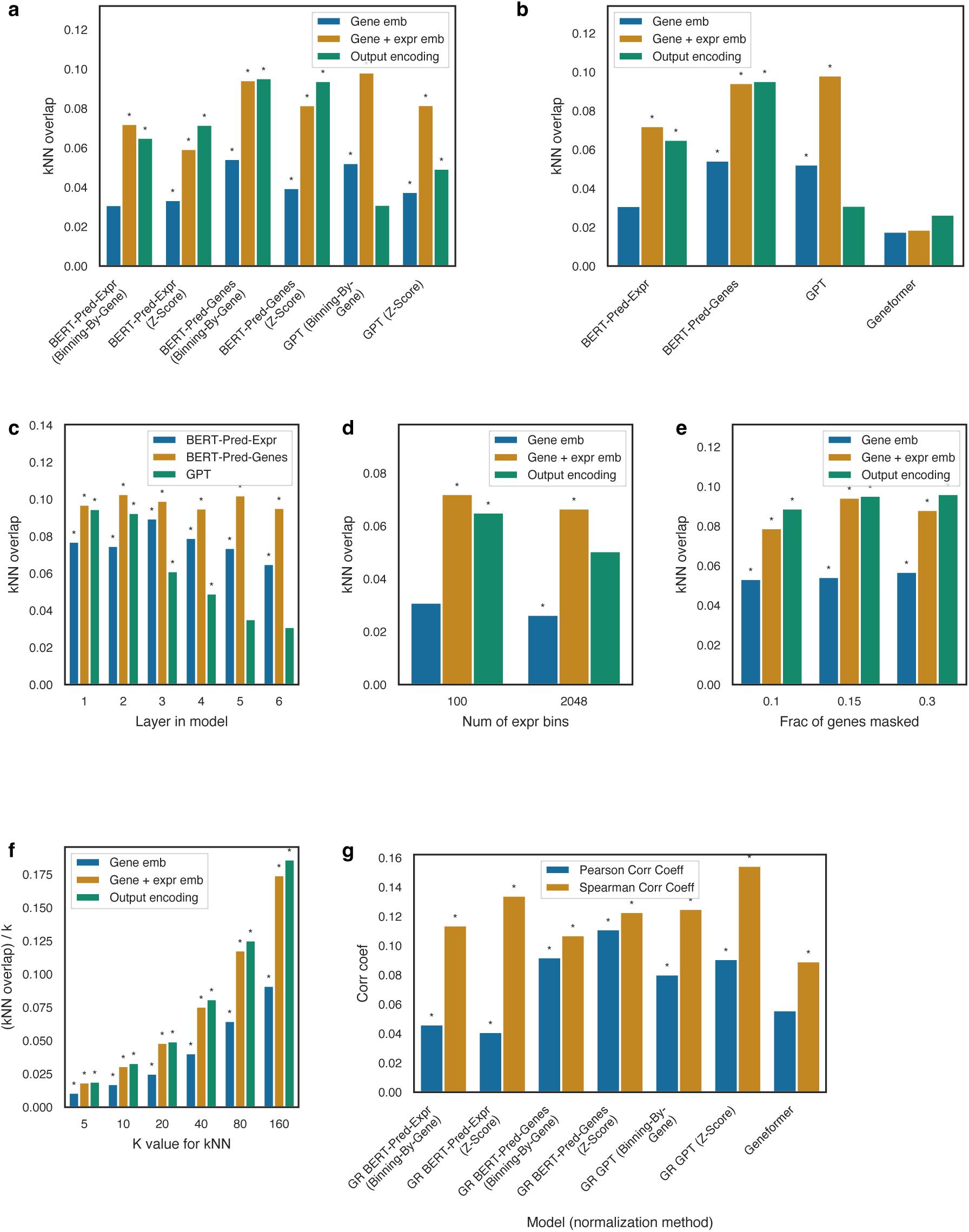
| Evaluating the capabilities of model encodings in mirroring genetic perturbation responses. This figure explores the abilities of various encodings – gene embeddings, gene plus expression embeddings, and hidden-layer encodings in the model – in reflecting transcriptomic changes due to genetic perturbations. This is quantified by the average observed overlap of the k (k=5 by default) closest neighbors in the model’s encoding space with k genes inducing most similar transcriptomic alterations. **a,** Observed overlap for various data normalization methods, **b,** Observed overlap in different GR and Geneformer models. **c,** Layer-specific observed overlap in different GR models. **d,** Observed overlap in BERT-Pred-Expr models with varying expression bin numbers. **e,** Observed overlap in BERT-Pred-Genes models with different fractions of genes masked. **f,** Distribution of ratios of overlap to the k value, across different k value for kNN analysis. **g,** Correlation coefficient between cosine similarities derived from cell embeddings *in silico* perturbations and those from actual transcriptomic responses in Perturb-seq experiments, across different models. * *p* < 0.01, as determined by a permutation test with 100 iterations.

We also examined each model’s ability in integrating gene expression information with gene identity information, by comparing gene + expression embeddings to gene only embeddings. We observed an increase in overlap after adding expression embeddings, indicating the models’ ability in incorporating the transcriptome state of cells to model gene regulatory correlations. In all three model architectures, normalization method of ‘Binning-By-Gene’ showed better performance than Z-Score-based method in these combined embeddings. In comparisons between models, our BERT-Pred-Genes and GPT models exhibited the best performance, with all of our models using ‘Binning-By-Gene’ normalization outperforming Geneformer (Fig. 4a, b and Supplementary Table 4).

We also evaluated the characteristics of encoding across different hidden layers within the models. It was found that BERT and GPT models exhibited divergent patterns, with BERT models showing stable overlap across deeper layers, while GPT models demonstrated a reduced representation of transcriptomic response information (Fig. 4c). Additionally, in the BERT-Pred-Expr model, increasing the number of expression bins led to a slight reduction in performance (Fig. 4d). However, the proportion of masked genes in BERT-Pred-Genes models did not obviously impact their ability to mimic transcriptomic responses to perturbations (Fig. 4e). Subsequent analysis of ratios of overlap to the k value in the kNN analysis revealed an increase in these ratios with larger k values (Fig. 4f). This trend suggests that the models are not only capturing local transcriptomic response relationships within a narrow scope of genes but are also effectively learning the global relationships among large number of genes. Lastly, comparison of observed overlap across epochs indicated that GPT models exhibited a faster learning rate compared to BERT models (Extended Data Fig. 8a-c).

### *In silico* perturbation experiment

We also assessed the model’s ability in simulating transcriptome responses to *in silico* gene knockdowns. Unlike previous evaluations focusing on static gene encoding, this analysis examined dynamic transcriptomic changes induced by *in silico* gene perturbations. In these experiments, we artificially altered the expression of specific genes and observed the resulting changes in representation of the entire transcriptome within the model. This is termed cell embedding, and is calculated by averaging the output layer encodings across all genes. The impact of gene knockdown on the transcriptome was quantified using cosine similarity between the baseline and perturbed cell embeddings. The *in silico* perturbation cosine similarity was then compared with the cosine similarity from observed transcriptome responses in Perturb-seq experiments.

Our experiments revealed that *in silico* knockdown of a gene within our model typically resulted in minimal changes to the cell embeddings (Extended Data Fig. 8d). This could be attributed to the dilution effect of perturbations of a single gene amidst the vast number of expressed genes in the bulk RNA-seq data, and our Transformer models allocate attentions to all the input genes. While the correlation coefficients were higher in our BERT-Pred-Genes and GPT models compared to Geneformer (Fig. 4g), further investigation should be helpful to illustrate the efficacy of this assessment, considering the minimal cell embedding changes in bulk RNA-seq samples.

### Using gene embeddings to train classifiers for biological feature predictions

We explored the utility of using the gene embeddings in our model, representing various biological attributes of genes across diverse dimensions, to extract different aspects of gene-related information and to perform different tasks. To facilitate information extraction and model training using a smaller number of samples, we further condensed the embedding to only 32 dimensions using Variational Autoencoder (VAE)^36^. We subsequently used the embeddings as features to train 5,050 classifiers (Methods), each of which was designed to predict a specific class in a biological attribute of genes, for instance if a specific gene is associated with a specific disease or not, or whether it is targeted by a specific transcription factor. Various categories of gene attributes were covered by the classifiers, including associations with rare and complex diseases, protein complexes, targeting by transcription factors, epigenetic modifications, glycosylation, Gene Ontology, protein domains, biological pathways, and cell type association (Extended Data Fig. 9a and Supplementary Table 5). The performance of a subset of exemplary classifiers is illustrated in Fig. 5.

**Fig. 5.**
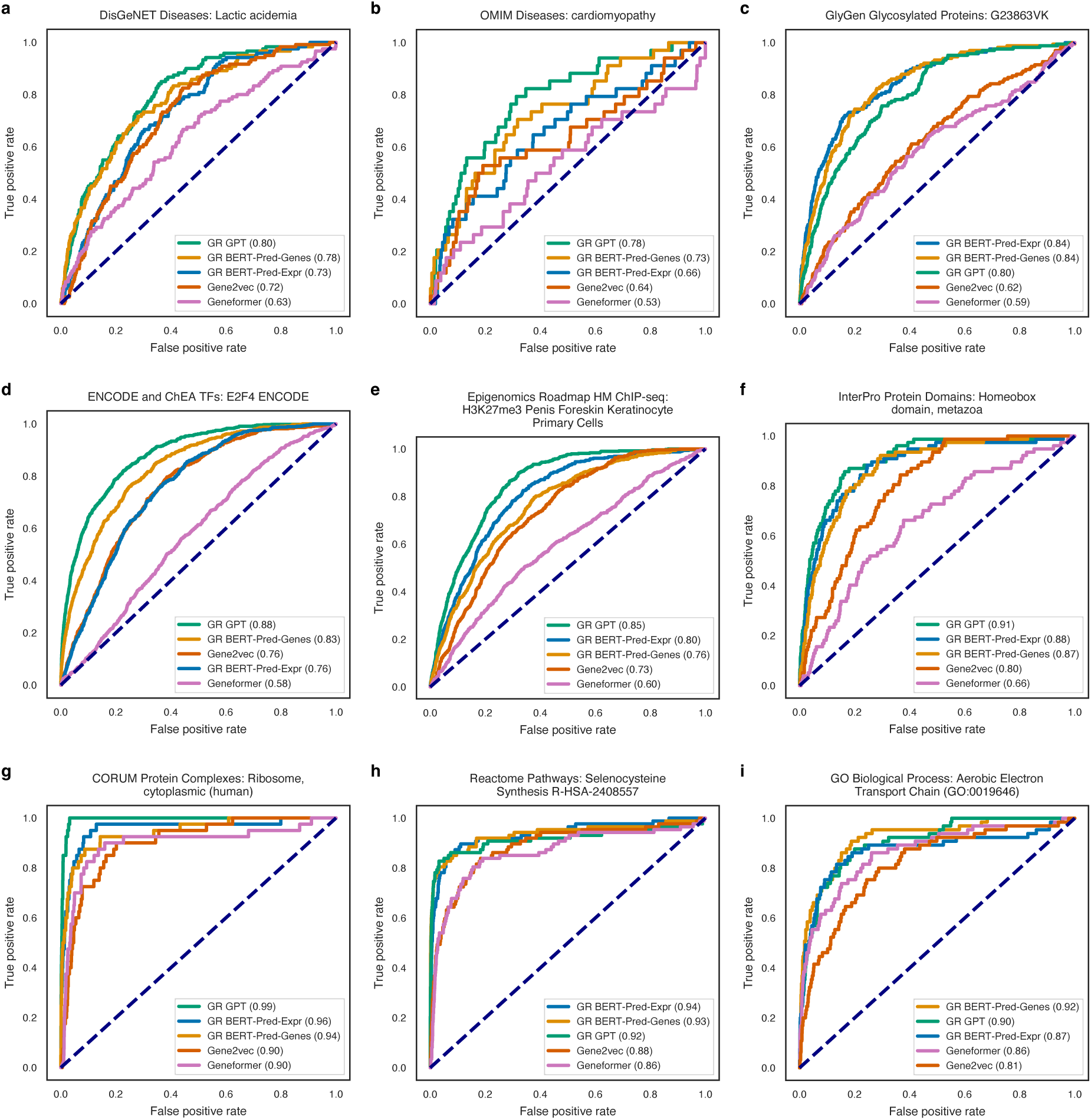
| Performance of gene attribute prediction classifiers. This figure demonstrates the capability of classifiers trained with gene embeddings as features to predict a specific class in some example gene attributes. Each ROC plot’s title indicates the database source and the specific gene class predicted by the classifier. **a,** ROC curves for classifiers differentiating lactic acidemia disease genes from non-lactic acidemia genes, based on data from the DisGeNet database. **b,** Classifier efficacy in distinguishing cardiomyopathy genes from non-cardiomyopathy genes, based on the OMIM database. **c,** Classifier performance in identifying genes associated with glycan G23863VK, referencing the GlyGen database. **d,** Performance of classifiers in predicting targets of the E2F4 transcription factor, utilizing data from the ENCODE database and ChEA data. **e,** Classifier accuracy in identifying genes marked by H3K27me3 histone modification in Penis Foreskin Keratinocyte Primary Cells. **f,** Classifier performance in predicting genes that encode proteins containing the homeobox domain. **g,** Efficacy of classifiers in identifying genes encoding proteins within the human cytoplasmic ribosome complex. **H** and **i,** Performance of classifiers in predicting genes involved in the Selenocysteine Synthesis pathway and Aerobic Electron Transport Chain gene ontology, respectively.

We then assessed the performance of models whose embeddings were used as features by examining the cumulative distribution of the Area Under the Curve (AUC) of all 5,050 classifiers. Compared to models using the Z-Score based normalization method, all models using the ‘Binning-By-Gene’ approach exhibited superior performance (Fig. 6a, Extended Data Fig. 10a, c and Supplementary Table 6-7). It was observed that all of our models that used the ‘Binning-By-Gene’ normalization significantly outperformed both Geneformer and Gene2vec (p < 1E-20, Kolmogorov-Smirnov test). Overall GPT showed the best performance (Fig. 6b, Extended Data Fig. 10b, d and Supplementary Table 6-7), with 72.65%, 34.67% and 14.20% of classifiers having AUC greater than 0.6, 0.7 and 0.8 respectively, compared to 20.36%, 11.60% and 8.20% for Geneformer. This indicates the superior effectiveness of the ‘Binning-By-Gene’ normalization method and the GPT model architecture in learning information for gene biological attribute prediction.

**Fig. 6.**
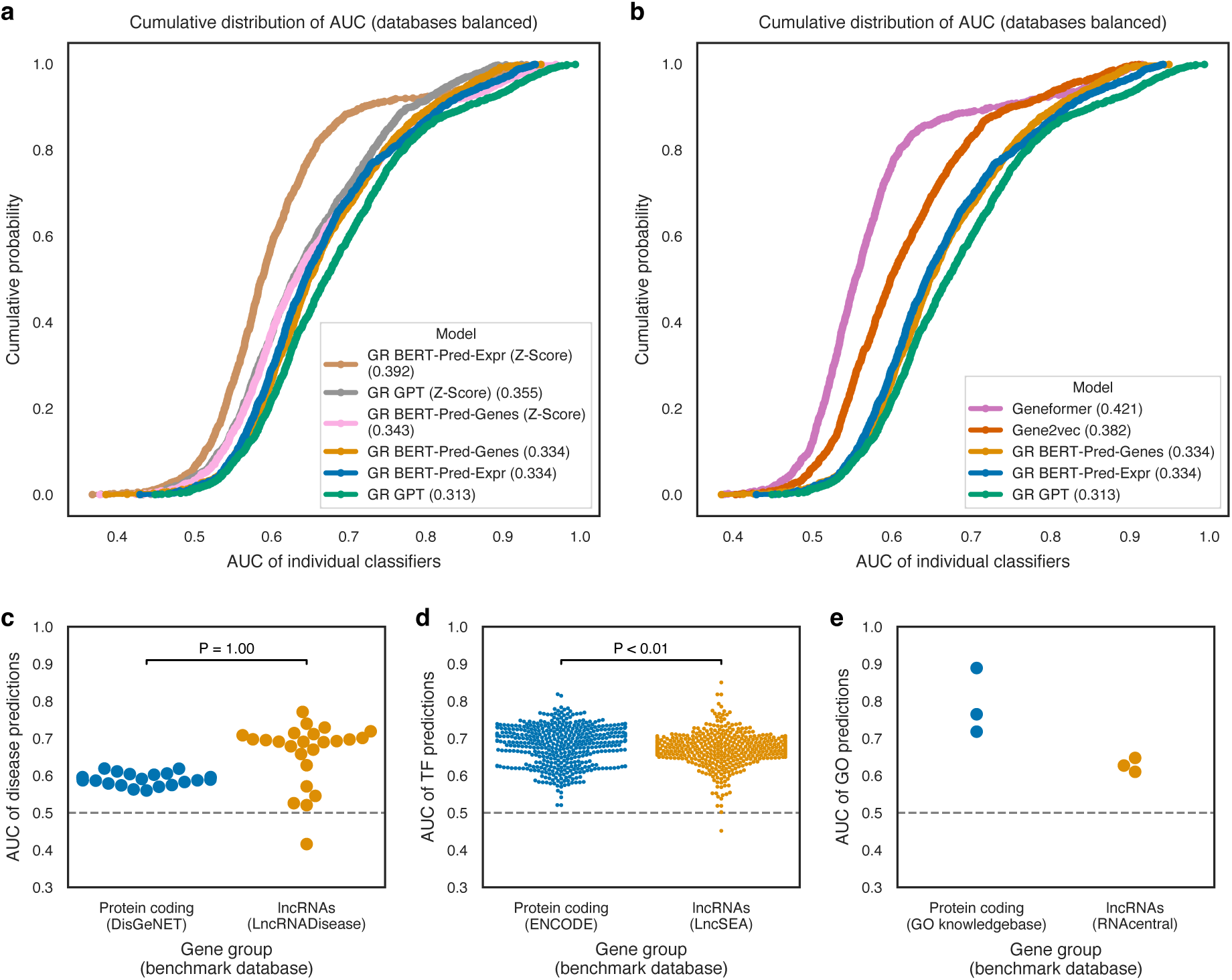
| Cumulative performance of all classifiers and evaluation on lncRNA attribute predictions. This figure provides a comprehensive view of the performance of all classifiers (with database balanced, each database contributed to a maximum of 100 classifiers) trained with gene embedding from different models. The plots display the cumulative distribution of the AUC values for these classifiers. The x-axis represents the AUC value, while the y-axis indicates the cumulative probability. Points on the plot represent the proportion of classifiers with AUC values less than or equal to the corresponding x-axis value. **a,** Cumulative distributions of AUCs for classifiers using encodings from models employing different normalization methods. The numbers in parentheses denote the area under the cumulative distribution curve, where lower values represent better classifier performance. In comparisons between Binning-By-Gene and Z-Score normalization, all pairs have p-values < 2E-4, two-sided Kolmogorov-Smirnov test. **b,** Cumulative distributions of AUCs for classifiers using gene embedding from GR models using Binning-By-Gene normalization methods, Geneformer and Gene2vec. For all pairs of comparisons, p-values are < 1E-4, except for GR BERT-Pred-Genes vs. BERT-Pred-Expr, which has a p-value of 0.15, two-sided Kolmogorov-Smirnov test. **c.** The distribution of AUC values for classifiers trained mostly on protein-coding genes, focused on predicting disease associations in both protein-coding genes and lncRNAs. Each dot represents one disease, with 24 diseases present in both benchmark databases being visualized. **d.** The distribution of AUC values for 457 classifiers trained mostly on protein-coding genes, on predicting targeted transcription factors in both protein-coding genes and lncRNAs. **e.** The distribution of AUC values for Gene Ontology (GO) predictions, where each dot represents a distinct GO. P-values are calculated using a one-sided Wilcoxon signed-rank test, aiming to determine if lncRNA AUCs are significantly lower than those of protein-coding genes. The p-value for GO predictions is not computed due to the limited sample size.

### Predicting biological attributes of lncRNAs

Our models learn information from the interrelationships between gene expressions, independent of the information of encoding proteins. This suggests that we can potentially make predictions without having the information of encoding proteins and make prediction on genes regardless of whether they encode protein or not. This approach opens up the possibility of applying classifiers trained on coding genes to predict biological attributes of long non-coding RNAs (lncRNAs)^37,38^, whose biological properties are less well characterized^39^. We undertook this analysis by training another GPT (’Binning-By-Gene’) model, using lncRNAs in addition to protein encoding genes as input. LncRNAs were therefore represented within the same embedding space as that of coding genes. We found that the model showed similar performance to the one trained on predominantly protein-coding gene data (Extended Data Fig. 9b-d). Following this, we employed the model’s embeddings to train 4,797 classifiers primarily on protein coding gene data (Methods, Supplementary Table 5), as previously described. Utilizing these classifiers, we made 62.5 million biological attribute predictions for 13,030 lncRNAs (Supplementary Table 8). These predictions included associations with 1,575 diseases/phenotypes, 1,007 Gene Ontologies, 477 pathways, associations with 469 cell types, and 1,268 transcription factor targets and epigenetic modifications.

To assess the transferability of classifiers trained mostly on protein-coding genes to lncRNAs, and to evaluate their performance, we used lncRNA-disease association data from the LncRNADisease database^40^ as a benchmark. There were 24 diseases with a sufficient number (n >= 30) of associated genes in both LncRNADisease database and DisGeNET^41^, the protein-coding gene disease database we used for training. For these 24 diseases, the classifiers’ performance on lncRNAs was found to be comparable to that on protein-coding genes (Fig. 6c and Supplementary Table 9).

Our model allowed us to predict disease-associated lncRNAs by transferring the knowledge of protein-coding genes. For example, we successfully predicted disease-associated lncRNAs discovered in recent years, including MIAT for atherosclerosis, MIR100HG, ELFN1-AS1, and LUCAT1 for colorectal cancer, HOTAIR and HNF1A-AS1 for esophageal carcinoma, and KCNQ1OT1 and H19 for myocardial infarction, among many others (Supplementary Table 10).

We further assessed classifier performance in predicting targeted transcription factors and GO terms^26,42^ for lncRNAs using LncSEA^43^ and RNAcentral^44^ databases as benchmarks, respectively. Evaluation of 457 classifiers for transcription factors revealed comparable effectiveness for both gene types (Fig. 6d and Supplementary Table 9). In the three GO categories with a sufficient number of lncRNAs (n >= 10) that can be predicted, the classifiers, despite some performance degradation, still demonstrated predictive capabilities beyond random chance (AUC = 0.5) (Fig. 6e and Supplementary Table 9).

## Discussion

In this study, we extensively evaluated Transformer-based self-supervised gene expression models for their ability in learning various biological attributes of genes. Additionally, we explored their capacity in simulating transcriptomic responses to perturbations by understanding gene relationships. We also explored strategies to enhance neural network training for learning the complex information within gene expression data. This involved evaluating various model architectures and configurations. As a result, we developed and implemented a novel normalization method with substantially improved performance, which entails imposition of a constraint where all genes have an equal chance to occupy every position in the model input. We demonstrated that, although the models were trained solely on gene expression data, they were able to learn a broad spectrum of biological information beyond gene expression. We subsequently explored the characteristics of various models, encodings in different layers, and performance in various tasks. We then demonstrated the versatility and capabilities of our approach by training 4,797 classifiers to make 62.5 million predictions on the biological attributes of 13,030 lncRNAs. We also performed validation that showcased the versatility and utility of our approaches.

The discovery that different biological attributes were learned by the models and represented by diverse dimensions in encoding is a promising finding. Our analyses indicated that, although trained with only gene expression data, the models have the capacity to learn information beyond expression. This outcome is reasonable, considering how the models learned to predict gene expression, and the fact that gene expression could be impacted by various biological factors. The process of back-propagation during training could guide the model toward learning any relevant information that could enhance the accuracy of expression prediction. The information of various gene attributes, such as associated pathways, transcription factor targeting, protein-protein interactions, and their correlation with gene expression, should be beneficial for improving the prediction. Thus, these attributes were learned by the models. This learned gene attribute information was captured in the gene embeddings, and the knowledge of how these attributes interact and correlate with gene expression was within the Transformer encoder’s parameters.

Of note was the observation that the GPT model demonstrated superior performance over the BERT models. The GPT approach has been only rarely used in gene network modelling^1–11^. GPT’s rapid learning curve, in comparison to BERT models, could be attributed to its architecture, which makes predictions for every gene in a transcriptome. This provides GPT with a variable and rich informational context for each prediction, enhancing its generalization capability by tackling different gene expression levels. In contrast, in each sample BERT models focus on predicting masked genes using information from unmasked ones, with a consistent informational context for each prediction. This could potentially increase the risk of relying on ‘shortcut’ information of co-expression^45^ from a few genes, despite the AdamW optimization method’s mechanism to penalize over-reliance on a limited number of features^46,47^. While BERT models demonstrate a slower learning rate than GPT, further investigation is required to illustrate the extent of additional information that can be acquired with extended learning periods.

In our study, we observed distinct characteristics within the encoding layers of BERT and GPT models. Notably, the representation of individual genes seems to diminish in deeper layers of GPT models, whereas it appears stable in BERT models. This phenomenon likely stems from the different ways information is processed by GPT and BERT architectures, reflecting how simple, isolated gene representations in lower layers are integrated into more complex and abstract events in deeper layers. Understanding these nuances could be beneficial for fine-tuning the model for specific tasks, guiding the selection of appropriate model architectures and layers for optimal information extraction.

The use of gene embeddings represents a new way of learning multiple gene attributes using only one type of omics data. By optimizing gene expression representation through innovative normalization methods and selecting suitable models and configurations, we can enhance our ability to learn gene attribute information. While further controlled comparisons and investigations would likely provide more evidence, the use of bulk RNA-seq data, with its more precise gene expression information, probably also contributed to improved model performance, despite sample size limitation.

The ability of these models to represent a wide range of biological attributes highlights their potential utility for numerous tasks beyond simply using them as features to predict gene attributes. By condensing this information into just 200, or even 32 dimensions, we could make these applications more feasible and attractive. This approach also shows promise as a means of overcoming the limitations associated with small sample sizes, as in many studies, large sample sizes are necessitated to overcome the loss in statistical power when conducting multiple tests across tens of thousands of genes.

It should be noted that the classification focuses on an aspect different from the clustering for the gene Attribute Learning Index. The clustering process was mainly used to show how well the model had learned biologically relevant information. For each gene attribute, clustering looked at all 200 gene embedding dimensions. However, typically, a gene attribute is relevant to only a subset of these dimensions. Consequently, the overall performance of the clustering might have been less than optimal, as the dimensions closely associated with a specific gene attribute were averaged with less relevant ones. The classifier training process could harness these associated dimensions and use them to make predictions.

In the study, we demonstrated one application of our gene representations. We used the knowledge of protein-coding genes to predict biological attributes of lncRNAs. Although our initial evaluations of the transferability of classifiers trained on protein-coding genes to lncRNAs yielded encouraging results, we acknowledge the potential influence of an RNA’s protein-coding status on the accuracy. With the availability of more extensive lncRNA annotation data, we will be able to undertake more thorough performance benchmarking. Nevertheless, our findings suggest that our approach represents a promising and potentially powerful methodology for predicting gene attributes, paving the way to more comprehensive insights into their roles in complex biological processes.

## Methods

### Preprocessing of Human Bulk RNA-seq Data

We obtained gene level human bulk RNA-seq data from the ARCHS4 database (version 2.2)^22^. All data collection and processing were conducted in compliance with ethical standards and procedures, with oversight and approval by the University of New South Wales (UNSW) Human Research Ethics Committee. We applied the following criteria for sample selection: total gene expression read counts exceeding one million, a minimum of 2000 genes with non-zero expression, and a single-cell probability below 0.5 (as calculated by the database) to exclude potential single-cell samples. These criteria were employed to facilitate the selection of high-quality samples. The samples were normalized by their library size to have a total read count of 10 million per sample. The read counts were then log-transformed (base 10). In handling duplicate gene symbols within the database, we retained only those with the highest mean expression. In benchmarking studies, for comparative purposes with Gene2vec^12^, our dataset was restricted to genes also present in Gene2vec, which includes protein coding genes and genes that encode non-coding RNAs. Furthermore, only genes with a mean normalized expression above 0.1 were maintained to exclude those with minimal expression across samples, resulting in a dataset of 17888 genes. In our lncRNA analysis, we also prepared a dataset with both protein coding genes and lncRNA genes included. In this dataset, all protein coding genes and lncRNA genes from ARCHS4 database with mean normalized expression > 0.1 were included, resulting in a dataset of 18739 protein coding genes and 13030 lncRNA genes. The normalized and filtered data were further normalized by one of the methods described below.

### Z-Score-based normalization

For each gene, we calculated the mean and standard deviation of its log-transformed expression values. Z-scores for each expression value were then computed using these parameters as follows:

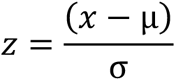

where x is the log-transformed expression value of a gene in a specific sample, μ and σ are the mean and standard deviation of the log-transformed expression values of the gene across all samples, respectively. We also explored the impact of including or excluding zero-expression genes in this calculation on model performance in the study.

### Binning-By-Gene normalization

Gene expressions across all samples were allocated into 2000 bins based on their expression rank, ensuring an equitable distribution. Specifically, the first bin was exclusively reserved for zero expression values. For non-zero expression values, we conducted a ranking process where each gene’s expression values were ranked across all samples. Subsequently, we adjusted the ranks of non-zero expression values by subtracting the count of zero expressions in each gene, ensuring that these adjusted ranks started from the lowest non-zero rank. The adjusted ranks were then used to distribute the non-zero expression values evenly across bins 2 to 2000. This method ensures that each bin, except the first, contains an approximately equal number of non-zero expression samples. The binning process was carefully designed to handle genes with varying numbers of zero expressions, thereby maintaining uniformity in bin sizes for non-zero expressions.

Pseudo code for Binning-By-Gene normalization:

*For each gene:*

1. *Rank all expression values (including zeros)*.
2. *Count the number of zero expressions for the gene*.
3. *Adjust the ranks for non-zero expressions by subtracting the count of zeros*.
4. *Calculate bin sizes based on the maximum adjusted rank:* *Bin size for non-zero expressions = max(adjusted rank) / (total bins – 1)*.
5. *Assign each non-zero expression to a bin:* *Bin = floor(adjusted rank / bin size) + 2*.
6. *Ensure that bin assignments do not exceed the total number of bins*.
7. *Assign zero expressions to the first bin*.

This process was repeated for all genes, ensuring that expressions were evenly distributed across bins, with the first bin reserved for zero expressions.

Prior to input into Transformer models, the gene expressions for each sample were ordered from highest to lowest based on z-scores or calculated bins (depending on the chosen normalization method). Only the top 2048 genes were inputted into models, attention was still applied on the zero expressions in the input if present. For the BERT-Pred-Expr model, expressions were allocated into either 2048 or 100 bins by their quantiles. The expression matrix was transformed into a float32 format, segmented into five sample-based chunks and saved as NumPy array files for more rapid loading and processing. Dataset was prepared using PyTorch’s Dataset class. By default, in BERT model, 15% of genes with non-zero expression had either their identity (gene symbol) or expression values masked. Gene embedding and expression embedding were added together in the BERT and GPT models.

### Preprocessing of Replogle et al. Perturb-seq data

The Perturb-seq dataset from the Replogle et al. study^29^ was obtained from the scPerturb database^48^. To align this dataset with our GeneRAIN models, which were pretrained on bulk RNA-seq transcriptomes, we generated pseudo-bulk RNA-seq data from the Perturb-seq dataset. This was achieved by averaging the single-cell transcriptomes from the same cell line and perturbation. These averaged transcriptomes were then normalized by library size and log-transformed (base 10). Z-scores were calculated using mean and standard deviation values for each gene derived from the ARCHS4 dataset. To apply the Binning-By-Gene normalization method on the perturb-seq dataset, bin boundaries were established based on the ARCHS4 dataset, with a subsampling to 0.5% of ARCHS4 samples to make data size more manageable. Same as the ARCHS4 data, for each sample, only the 2,048 highest expressed genes were inputted into BERT or GPT models.

Control samples from the database, representing cell lines without any genetic perturbation, were designated as baseline samples for our downstream analysis. For each genetic perturbation in the dataset, we randomly selected 10 control samples from the same cell line to represent the baseline transcriptome states before the perturbation. The resulting post-perturbation transcriptome response was represented by the observed gene expression in the Perturb-seq experiment.

Each entry in the resulting dataset comprised the baseline expression profile, a single perturbed gene, and the post-perturbation expression profile. For *in silico* gene knockdown experiments, the perturbed gene was assigned the lowest expression bin among the genes inputted into the model. This also means that *in silico* knockdown impacts only the 2,048 highest expressed genes which are selected for model input.

### Model hyper-parameters

We used models of BertForMaskedLM and GPT2LMHeadModel from Hugging Face’s transformers library^49^. In our models, each gene and expression value was represented in a 200-dimensional space. The model architecture is composed of six layers, each with four attention heads, and each head has 32 dimensions. We used embedding dropout^50^ rate of 0, attention dropout rate of 0.05 and feed-forward network dropout rate of 0.05. The epsilon value for layer normalization^51^ was set at 1e-10. The models employ the Gaussian Error Linear Unit (GELU)^52^ as the activation function in its hidden layers. We adopted a ‘norm-first’ strategy, applying layer normalization before other operations within each transformer block.

### Model training

For efficient data processing across multiple GPUs, we utilized the Distributed Data Parallel (DDP) framework. For optimization, the AdamW optimizer^46,47^ was employed, with a base learning rate of 0.00001 and a maximum learning rate of 0.0001. A weight decay of 0.01 was applied to regularize the learning process, and the initial learning rate was set to 0.00002, which represents one-fifth of the maximum rate. The learning rate was modulated using the OneCycleLR scheduler during the first 30 epochs, followed by a constant learning rate equivalent to that of the 30th epoch for subsequent epochs. The initial phase of the learning rate increase was set at 20% of total training iterations, with a division factor of 5. The total steps for the scheduler were set at 40, and the number of epochs was set at 40, and the step size up for learning rate increase was configured at 4. The models after 40 epochs of training were used for downstream tasks by default.

To balance memory usage and batch size, we set gradient accumulation steps at 5. The batch size was set to be 12. The data was partitioned into training and validation sets, allocating 90% of the dataset for training.

### Analysis of gene attribute learning in gene embedding

We sourced gene attribute databases from the Enrichr database^53–55^, as detailed in Supplementary Table 3 ^26,42,56–73^. In this analysis, we evaluated how much biological attribute information of genes was learned in gene embeddings by the models. This was achieved by interrogating the performance of unsupervised clustering using gene embeddings in reconstructing the biological grouping relationships recorded in these gene attribute databases.

To ensure clarity in gene membership, for each biological attribute from a database (e.g. disease association from ClinVar database^66^), we first eliminated genes assigned to two or more groups, such as genes that are associated with multiple diseases. We focused exclusively on genes with unique group memberships. Furthermore, for each comparison between two models (e.g., GR-GPT vs. Geneformer), we filtered the genes to include only those present in both models, ensuring consistent gene sets were used for comparative analysis. Additionally, we retained only gene groups comprising a minimum of five genes, enabling robust clustering operations.

In each clustering experiment, we randomly selected four gene groups (e.g. groups of genes associated with disease A, B, C and D) from the target database and performed K-means clustering using the gene embeddings from the models under comparison. To assess the clustering performance, we calculated three metrics, including the Adjusted Rand Index (ARI)^24,25^, Normalized Mutual Information (NMI)^28^, and Fowlkes-Mallows Index (FMI)^27^, using functions of adjusted_rand_score, normalized_mutual_info_score and fowlkes_mallows_score from python package sklearn.metrics, respectively. ARI measures the similarity between two clusterings, considering all pairs of samples and counting pairs that are assigned in the same or different clusters in the predicted and true clusterings. FMI evaluates the geometric mean of the precision and recall of the clustering, providing insight into the accuracy and robustness of the clustering. NMI is a Normalization of the Mutual Information score that measures the shared information between two clusterings, adjusted for chance. These metrics collectively offer a comprehensive view of the ability of the embeddings to replicate the biological groupings. The clustering experiment was repeated 100 times. We compared these results with a null distribution derived from 100 experiments of random permutations of the gene embeddings, to evaluate the significance of the observed clustering metrics values. The gene Attribute Learning Index is thus the average difference between the actual metric values and those from random permutations, across ARI, NMI, and FMI, over the 100 experiments.

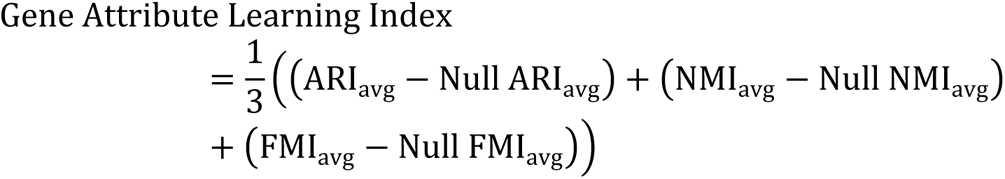

with:

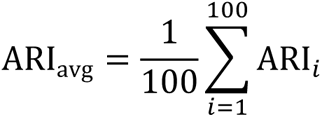

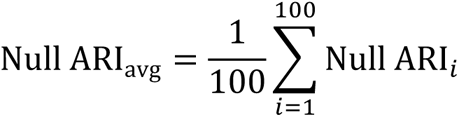

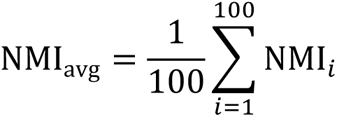

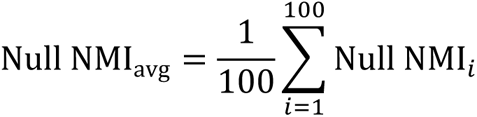

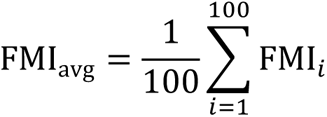

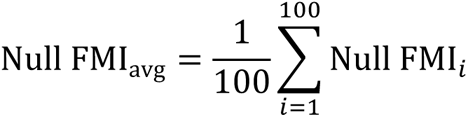

where ARI_avg_, NMI_avg_ and FMI_avg_ are the average ARI, NMI and FMI calculated over 100 iterations, respectively, and Null ARI_avg_, Null NMI_avg_ and Null FMI_avg_ are the average ARI, NMI and FMI calculated over 100 iterations with random permutations of the gene embeddings, respectively.

To assess the importance of each embedding dimension in gene clustering, we computed the F-statistic for each dimension using Analysis of Variance (ANOVA). This process enabled the quantification of the variance each dimension contributed to separating gene clusters. Subsequently, these F-statistics were normalized across all dimensions, to make the normalized F-statistics across all dimensions have a sum of 1.0. Dimensional importance for each embedding was accumulated across 100 clustering experiments. This approach facilitated a more interpretable and comparative representation of how each dimension in the embedding space contributed to the differentiation of gene clusters.

### Evaluation of model encoding using Perturb-seq data

In this analysis, we aimed to explore whether gene encoding within our model encapsulated information about the transcriptome’s response to gene perturbation. We investigated by checking if genes causing similar transcriptomic responses have similar encodings. Additionally, we examined if integrating expression embedding (representing transcriptome context) improves this transcriptomic response information. A comparative analysis across different model layers and between various models was also conducted.

To extract the encoding, we fed the transcriptome of interest into the models. For our GR models, we used gene expression data from 100 pseudo-bulk RNA-seq control samples prepared above from the Replogle et al. study^29^. The mean encoding of each gene across these samples was computed at the layer of interest and used as the gene’s representative encoding. For Geneformer, which was trained using single-cell RNA-seq data, gene expression data from 500 single-cell control cells from the Replogle et al. study was used. This data was preprocessed using the functions from the Geneformer Hugging Face repository, with gene encoding subsequently averaged across the 500 cells. Samples and cell lines from ‘K562_essential’ experiment in the study were used for both our models and Geneformer. Extraction of the combined gene and expression embedding and output encoding from each hidden layer was achieved by enabling the ‘output_hidden_states’ parameter in the model’s ‘forward’ function.

For comparison of encodings between two different models (e.g., GR-GPT vs. Geneformer), to avoid potential bias caused by the difference in the genes included by the models, we confined our analysis to genes present in both models. This approach ensured a consistent gene set for comparative evaluation. Utilizing the normalized expression data from Perturb-seq, we identified genes with the most similar perturbation response using the k-Nearest Neighbors (kNN) overlap method, with k=5 by default. We also determined the closest neighbors in the model encoding space. The degree of overlap between these two sets of neighbors was calculated for each gene, providing a measure of the encoding’s ability in mirroring the transcriptomic response to perturbations.

To measure the significance of the observed overlap, we employed a permutation test. In this test, one space was randomly shuffled 100 times by default, and the overlap was calculated to generate a distribution of overlaps under the null hypothesis of no association between the two spaces. The p-value was determined by the proportion of random overlaps greater than or equal to the observed overlap.

### *In silico* gene perturbation

This analysis aimed to assess the model’s ability in simulating transcriptome responses to *in silico* gene knockdowns. Unlike previous evaluations focusing on static gene encoding, this analysis interrogated dynamic transcriptomic changes induced by *in silico* gene perturbations.

For our GR models, we randomly selected 10,000 perturbation samples from the Perturb-seq dataset. Each sample comprised a baseline expression profile alongside a perturbed profile, in which only the expression of the targeted gene was altered, keeping expressions of other genes identical to the baseline. The baseline expression profile and perturbed profile were fed into the model separately, and the resulting encodings in the output layer of the model were extracted. The difference between output encoding from the baseline and perturbed expression profiles were viewed as the transcriptomic response predicted by the model.

We focused on the model output from the final layer, considering it embodies the most abstract and comprehensive representation learned by the model. As the output layer contains encoding for every gene, to get a single representation of cell transcriptome state, we calculated the average embedding across genes to derive a single cell embedding, representing the transcriptomic state of the cell. To quantify the extent of transcriptomic response as predicted by the model, we computed cosine similarities between the baseline and perturbed cell embeddings. Correspondingly, cosine similarities were calculated using actual RNA-seq results from the Perturb-seq experiment.

The agreement between model-predicted and actual cosine similarities was quantified using Pearson and Spearman correlation coefficients. This statistical approach enabled us to evaluate the model’s accuracy in mimicking transcriptomic responses to genetic perturbations.

In the case of Geneformer, a single-cell RNA sequencing model, we chose 300 control cells from the Perturb-seq study as baseline references. For each cell, every gene with non-zero expression was subjected to an *in silico* knockdown, treating each knockdown as a separate perturbed cell instance. Samples and cell lines from the ‘K562_essential’ experiment were consistently employed for both our models and Geneformer.

### Training classifiers using gene embeddings from AI models

This analysis aimed to assess the informational content within the gene embeddings generated by AI models. Specifically, we focused on their utility in predicting various biological attributes of genes, such as their association with specific diseases. We trained a total of 5,050 classifiers, each designed to predict a specific class in a biological attribute of genes, using gene embeddings as features. For instance, for the biological attribute of disease association, the trained classifier was to predict if a gene is associated with a specific disease or not. To ensure consistency in comparisons, we utilized only those genes that were represented across all the studied models.

Before classifier training, we compressed the gene embeddings into a more compact representation using Variational Autoencoders (VAEs)^36^. This dimensionality reduction facilitated the training with a limited number of gene examples. The encoder in VAE had a linear transformation layer with ReLU activation^74^ reduced the input dimensions to a 32-dimensional latent space. This latent space was characterized by two parameters, mean (μ) and log variance (log σ²), calculated through distinct linear layers. The decoder in VAE used a linear layer to reconstruct the gene embeddings from the latent space.

During training, the training dataset was partitioned, allocating 10% for validation purposes. The VAE underwent training over 2000 epochs, utilizing the Adam optimizer^47^ with a learning rate set at 1e-4. The loss function incorporated mean squared error (MSE) for evaluating reconstruction accuracy, along with Kullback-Leibler (KL) divergence^75^ for latent space regularization. This combination ensured both the accuracy of reconstructed data and the effectiveness of the dimensionality reduction process. A separate VAE was trained for gene embedding from each of the studied models.

After dimension reduction, to assess the predictive power of gene embeddings on biological attributes, we trained 5,050 distinct classifiers. Each classifier was dedicated to predicting a specific class within a gene attribute. The training data for these classifiers were obtained from Enrichr^53–55^, except for Neuromuscular disease (NMD) genes data, which was obtained from https://www.musclegenetable.fr, as detailed in Supplementary Table 5 ^26,41,42,56,58,63,65,66,69,70,72,76–80^. For NMD genes, genes known to cause monogenic congenital muscular dystrophies, congenital myopathies, distal myopathies and other myopathies in the database were categorized as ‘certain skeletal muscle diseases’ to reflect their more certain association with skeletal muscle disorders. Furthermore, this category, along with genes known to cause muscular dystrophies, myotonic syndromes, ion channel muscle diseases, malignant hyperthermia and metabolic myopathies were collectively categorized as ‘all skeletal muscle diseases’.

For each class within a database (e.g., a specific disease in the ClinVar disease database^66^), a classifier was trained to predict the association of genes with that class. After intersecting genes from the database with genes available in model embeddings, classes from databases were filtered to exclude those with only small number of genes available, based on criteria detailed in Supplementary Table 5. The classifiers utilized gene embeddings as features and corresponding biological attributes as labels. By default, classification aimed to distinguish between genes associated with a target class (e.g., a specific disease) and all other genes. We also experimented the classification aim of distinguishing between genes associated with a target class and all other classes in the same database. We employed a 5-Fold cross-validation method, dividing the dataset into five parts, each used once as a test set. During each iteration, only the training set was utilized for model training, with no exposure to the test set for parameter tuning.

There was class imbalance in the dataset (e.g. fewer genes associated with a specific disease compared to non-associated genes), and we applied a Random UnderSampler using imblearn python library to balance the classes by subsampling the major class. This approach ensured diversity in resampling by varying the random state in each iteration. To overcome information loss from major class subsampling, we adopted an ensemble classifier strategy^81^. This involved training five individual classifiers, with each classifier trained on uniquely subsampled data. The ensemble’s final prediction was an average of the probabilities predicted by the individual classifiers. The ensemble’s performance was evaluated on the test set, unseen by any individual classifier. Two classification methodologies were tested: Linear Discriminant Analysis (LDA)^82^ and XGBoost (XGB)^83^. The results used for visualization were from LDA by default.

To evaluate and visualize the overall performance of all classifiers using the gene embedding of a model of interest features, the cumulative distribution of the AUC values for all the classifiers was plotted. To prevent overrepresentation by classifiers from databases with a larger number of gene classes, we also created a cumulative distribution plot from database balanced classifiers. In this balancing process, classifiers from databases with more than 100 gene classes were down-sampled to 100. This strategy ensured that the resulting curve more accurately reflected the model encoding’s ability to predict a diverse range of gene attributes.

### Predicting biological attributes of lncRNAs

For this task, GR GPT model was trained on the dataset of 18,739 protein coding genes and 13,030 lncRNA genes prepared previously, using the ‘Binning-By-Gene’ method for normalization. For classifier training, to facilitate comparison with previously trained classifiers, genes were filtered to include only those also present in Geneformer. This resulted in a gene set of 18,358 protein-coding genes and 353 lncRNA genes. The embeddings of these genes in the GPT model were used as features to train classifiers, and to predict the gene attributes listed in Supplementary Table 5. We excluded databases that describe protein domains, protein complexes, and protein glycosylation status, as these are not relevant to lncRNAs. The trained classifiers were then utilized to predict the lncRNA attributes using their embeddings. Prediction probabilities were averaged across five classifiers, each from one iteration of 5-fold cross-validation process.

For evaluating classifier performance on lncRNAs, we first evaluated the performance in predicting disease associations. Data from the file ‘website_causal_data.tsv’, obtained from the LncRNADisease v3.0 database^40^ was used as a benchmark. For more accurate AUC estimation, disease filtering was conducted to retain only those with >=30 associated lncRNAs covered by our GR model, with a total of 24 diseases remained for evaluation. In this process, the classifiers were assessed based on their ability to distinguish between lncRNAs associated with a target class (in this evaluation, a specific disease) and all other lncRNAs covered by our GR model, or similarly to distinguish between protein coding genes associated with a target class and all other protein coding genes covered by our model.

To evaluate the performance of targeted transcription factor prediction, classifiers primarily trained on protein coding genes from ENCODE_TF_ChIP-seq_2015 dataset of Enrichr database were examined. Data from the file of ‘set_Transcription_Factor.txt’ downloaded from the LncSEA 2.0 database^43^ were used as a benchmark, transcription factors marked as targeting the ‘Promoter_2kb_1kb’ region of lncRNAs were used. For more accurate AUC estimation, transcription factor filtering was conducted to retain only those with >=30 associated lncRNAs covered by our GR model. A total of 457 transcription factors remained for evaluation.

For evaluating performance of GO predictions, data labeled ‘rnacentral_rfam_annotations’ from the RNAcentral database^44^ were used. Here, lncRNA IDs were converted into gene symbols or Ensembl IDs utilizing the ‘id_mapping.tsv’ file from the same database. Due to the limited number of GO terms with sufficient number of associated lncRNAs, the filtering criteria were relaxed to include GOs >=10 associated lncRNAs. Ultimately, 3 GO terms met these criteria and were used for evaluation.

In all evaluations of classifier performance on lncRNAs, genes used in the training dataset were excluded to avoid biases in the assessment.

## Data availability

The data used to train GeneRAIN and the results it produced are accessible online at https://zenodo.org/records/10408775. This includes the ‘human_gene*.npy’ files, which are normalized datasets from human bulk RNA sequencing (ARCHS4) used for training. The subsampled dataset which can be used for determining binning boundaries in Binning-By-Gene normalization of new data is available in data.tar.gz. The 200-dimensional and 32-dimensional gene representations, namely GeneRAIN-vec.200d and GeneRAIN-vec.32d, derived from the GR GPT protein-coding+lncRNA model, are also available. The prediction results of all gene attributes for all protein-coding and lncRNA genes from GR GPT model can also be found in ‘data.tar.gz’. For more information on how to use this dataset, please see the README file at https://github.com/suzheng/GeneRAIN/blob/main/data/README.md. Provided files are made available under the Creative Commons Attribution Non Commercial 4.0 International License. All data is free to use for non-commercial purposes.

## Code availability

The checkpoints for the GeneRAIN protein-coding gene pretrained models (including BERT-Pred-Genes, BERT-Pred-Expr and GPT models) using Binning-By-Gene normalization, as well as the GeneRAIN GPT protein-coding+lncRNA pretrained model are available in file ‘data.tar.gz’ at https://zenodo.org/records/10408775. The source code for tokenizing transcriptome data, training the GR models and preparing datasets for new data using the Binning-By-Gene normalization method is available at https://github.com/suzheng/GeneRAIN. Provided code is made available under the Creative Commons Attribution Non Commercial 4.0 International License. All code is free to use for non-commercial purposes.

## Acknowledgements

We would like to thank the Research Technology Services (ResTech) at the University of New South Wales for their help with computational resources, with a special thank you to Martin Thompson for his support and assistance. We also extend our thanks to the National Computational Infrastructure (NCI) for their support. This research was undertaken with the assistance of resources and services from the National Computational Infrastructure (NCI Australia), which is supported by the Australian Government through the National Collaborative Research Infrastructure Strategy (NCRIS). Z.S. acknowledges funding support from The University of New South Wales (University Postgraduate Award scholarship).

## Author contributions

Z.S. conceptualized and designed the research, as well as developed GeneRAIN. E.O., M.D. and F.V. provided project supervision and strategic guidance on research methodologies and analysis. Z.S. and M.F. conducted data analysis, generated visualizations, and led the writing of the manuscript. All authors contributed to the intellectual content, provided critical feedback, participated in revising the manuscript, and approved the final version for publication.

## Additional information

Supplementary Information is available for this paper.

## Extended Data

**Extended Data Fig. 1.**
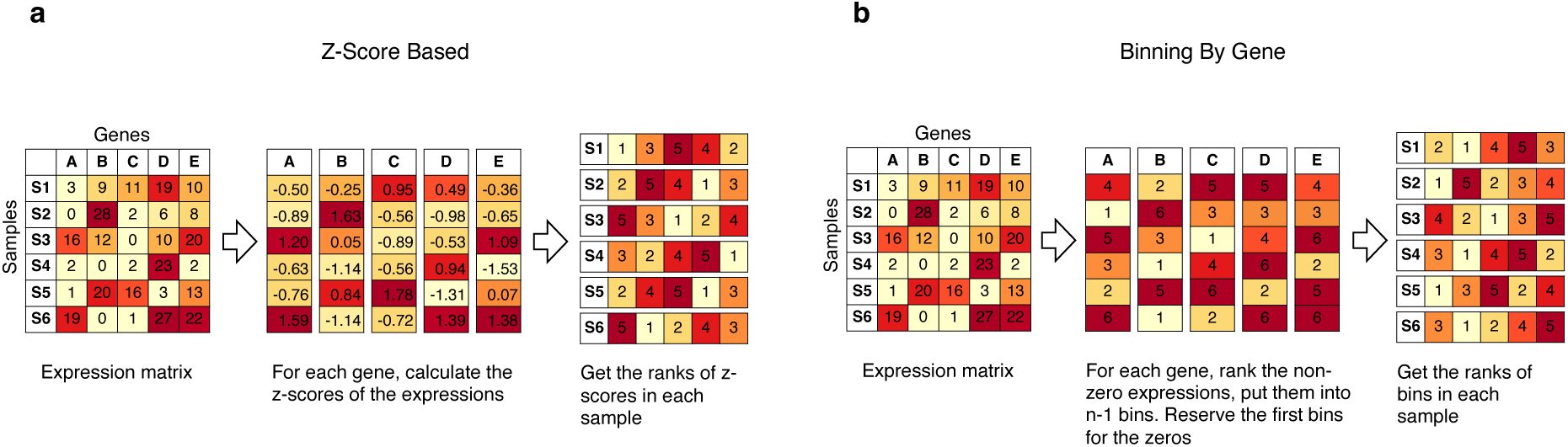
| Illustration of two normalization methods. **a**, Z-Score-based normalization: The left panel depicts a matrix of sample-by-gene expression levels, with genes represented by letters (A, B, C, etc.). For each gene, the mean and standard deviation of its expression across samples are calculated to derive the z-scores (middle panel). The right panel demonstrates how these z-scores are used to rank genes within each sample, which are then used for input to the BERT or GPT models. **b,** Binning-By-Gene normalization: This approach involves ranking expressions across samples for each gene and allocating them into specified bins (n=6 in this example) based on quantile ranks (middle panel). The lowest bin is reserved for genes with zero expression. Subsequently, these bin allocations are used to rank the genes within each sample, which are then used for input to the BERT or GPT models. In this illustrative example, there are instances of genes within the same sample have identical bins (or tied bins), they are randomly assigned neighboring ranks.

**Extended Data Fig. 2.**
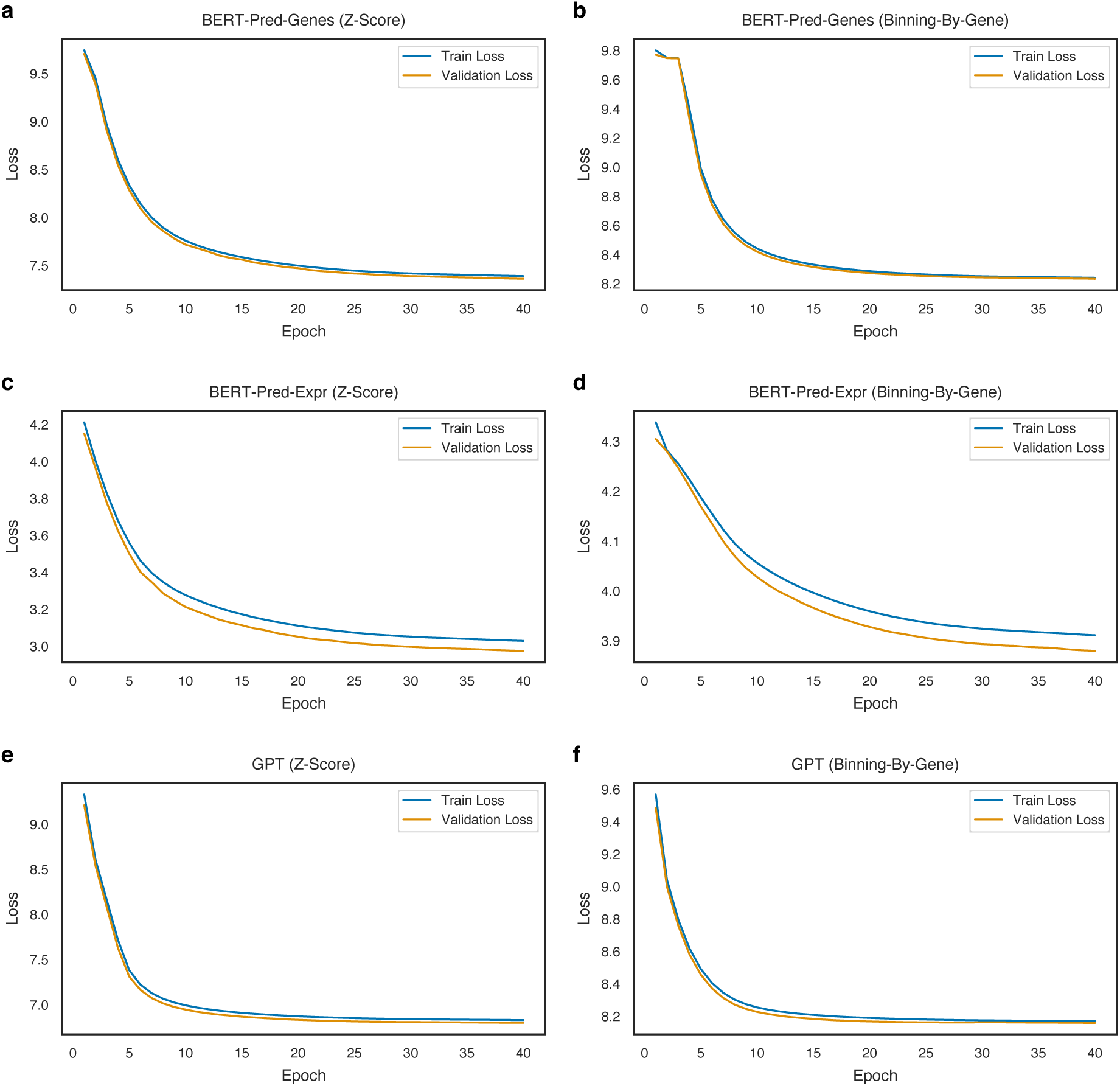
| Training and validation loss across epochs. This figure illustrates the training and validation loss over epochs for various models, including BERT-Pred-Genes, BERT-Pred-Expr, and GPT, each of which employs two normalization methods: Z-Score-based and Binning-By-Gene.

**Extended Data Fig. 3.**
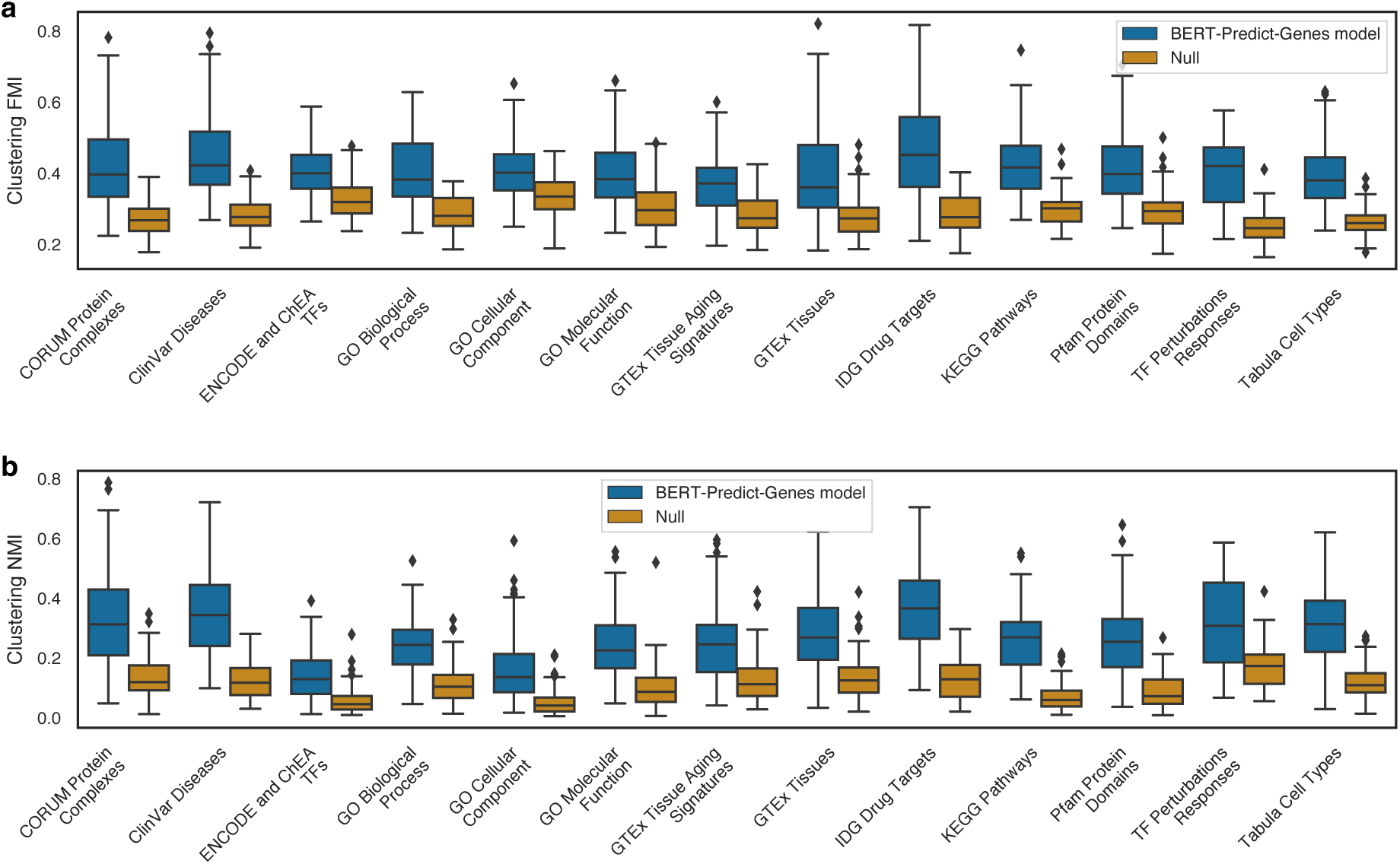
| Clustering agreement of gene embeddings from BERT-Predict-Genes model. This boxplot illustrates the clustering agreement of gene embeddings, as learned by our BERT-Predict-Genes model, with grouping of genes within various databases. The agreement is measured using **a,** the Fowlkes-Mallows Index (FMI) and **b,** Normalized Mutual Information (NMI), compared against a null permutation distribution. The consistency of clustering is determined based on 100 random selections of four gene groups from each database. In each boxplot, center line denotes the median; box limits represent the upper and lower quartiles; whiskers extend to 1.5× interquartile range; outliers are depicted as points. In all comparisons, p-values were < 1E-9, determined by a two-sided Mann-Whitney U test with Benjamini-Hochberg adjustment.

**Extended Data Fig. 4.**
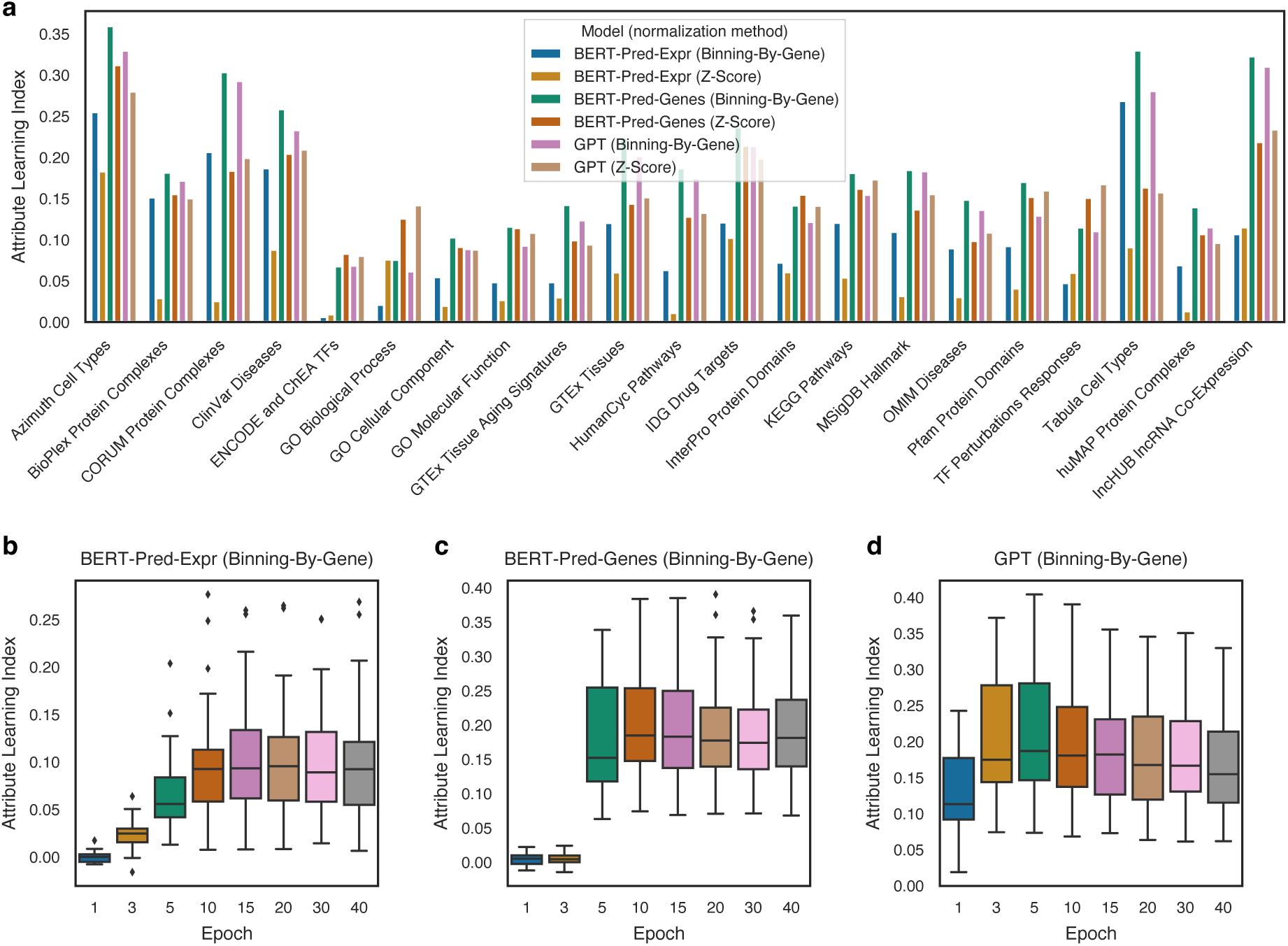
| Breakdown of gene Attribute Learning Indices by database and training epoch. **a**, Detailed breakdown of gene Attribute Learning Indices by database for models using different normalization methods. **b,** Distribution of gene Attribute Learning Indices over epochs for the BERT-Pred-Expr model, **c,** the BERT-Pred-Genes model, and **d,** the GPT model. Each box in boxplots encapsulates 21 data points, representing 21 different databases of gene attributes. Center line, median; box limits, upper and lower quartiles; whiskers, 1.5× interquartile range; outliers, points.

**Extended Data Fig. 5.**
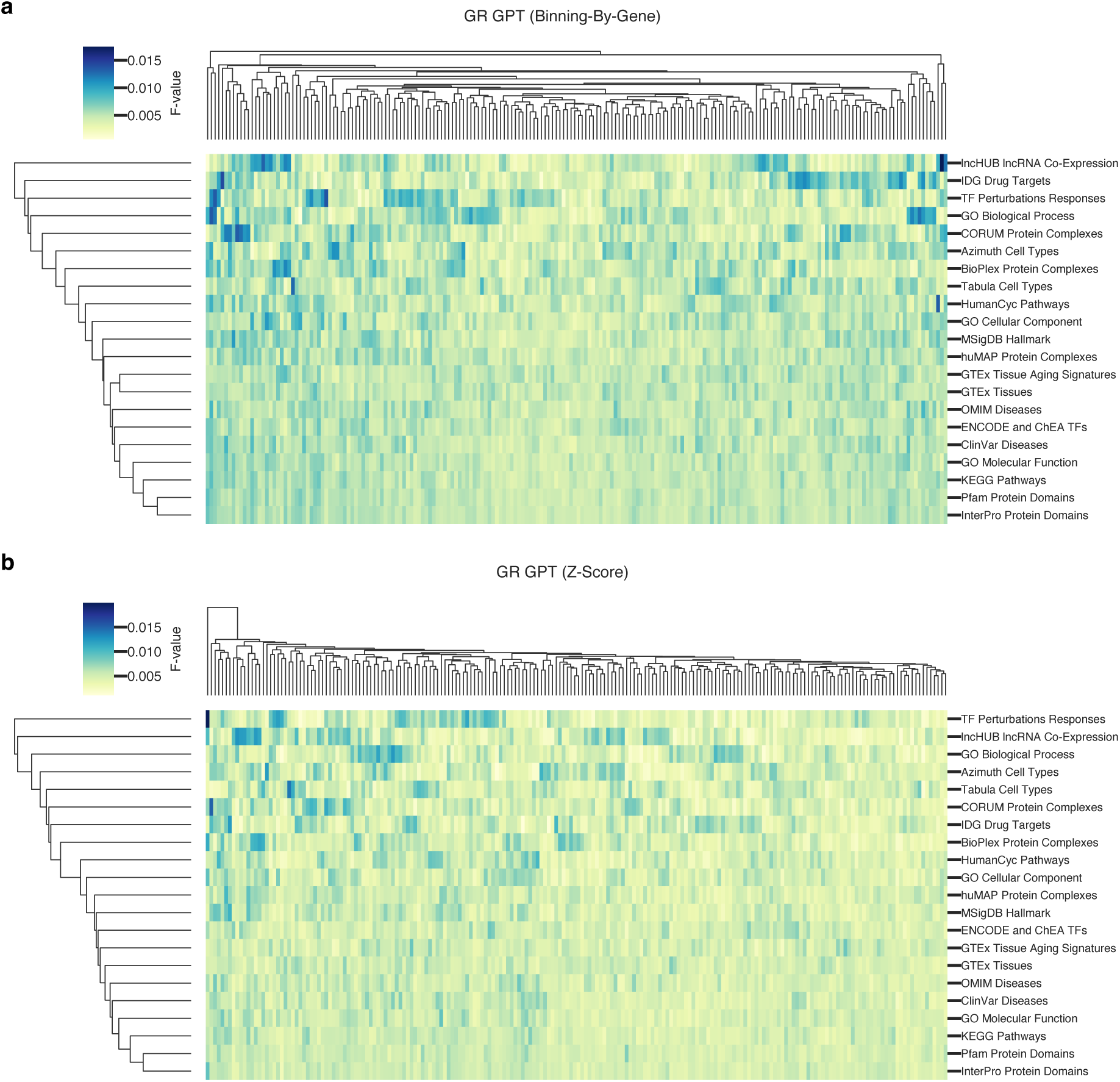
| Dimensional contributions in gene embeddings to gene attribute learning. This figure depicts the relative importance of each embedding dimension in clustering biological attributes of genes, as indicated by the color intensity representing the normalized F-statistic from ANOVA. **a,** Contribution distribution in the GPT model using Binning-By-Gene normalization, with each row corresponding to gene attributes from a specific database and columns denoting embedding dimensions. **b,** The distribution for the GPT model using Z-Score-based normalization.

**Extended Data Fig. 6.**
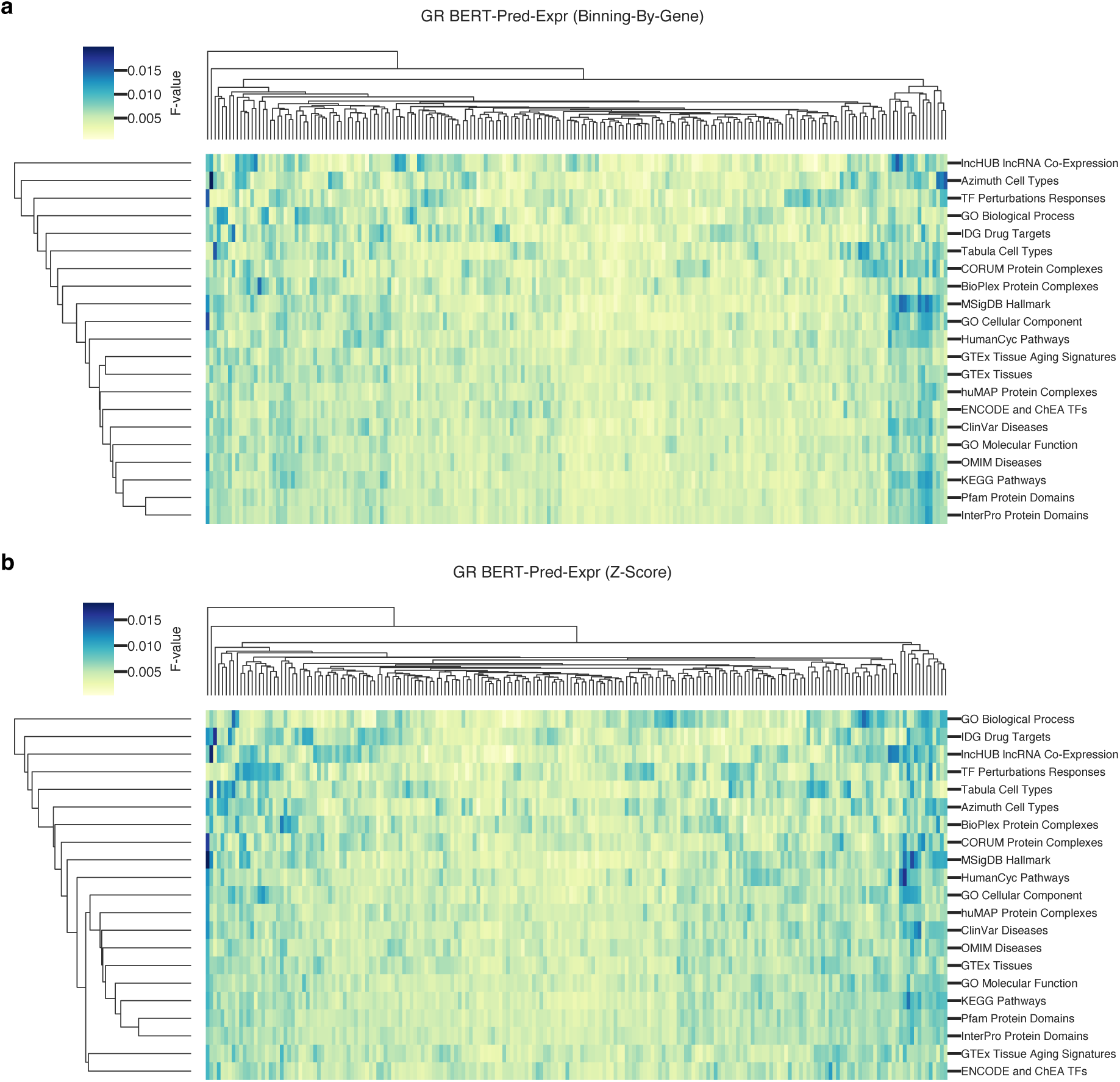
| Dimensional contributions in gene embeddings to gene attribute learning. This figure depicts the relative importance of each embedding dimension in clustering biological attributes of genes, as indicated by the color intensity representing the normalized F-statistic from ANOVA. **a,** Contribution distribution in the BERT-Pred-Expr model using Binning-By-Gene normalization, with each row corresponding to gene attributes from a specific database and columns denoting embedding dimensions. **b,** The distribution for the BERT-Pred-Expr model using Z-Score-based normalization.

**Extended Data Fig. 7.**
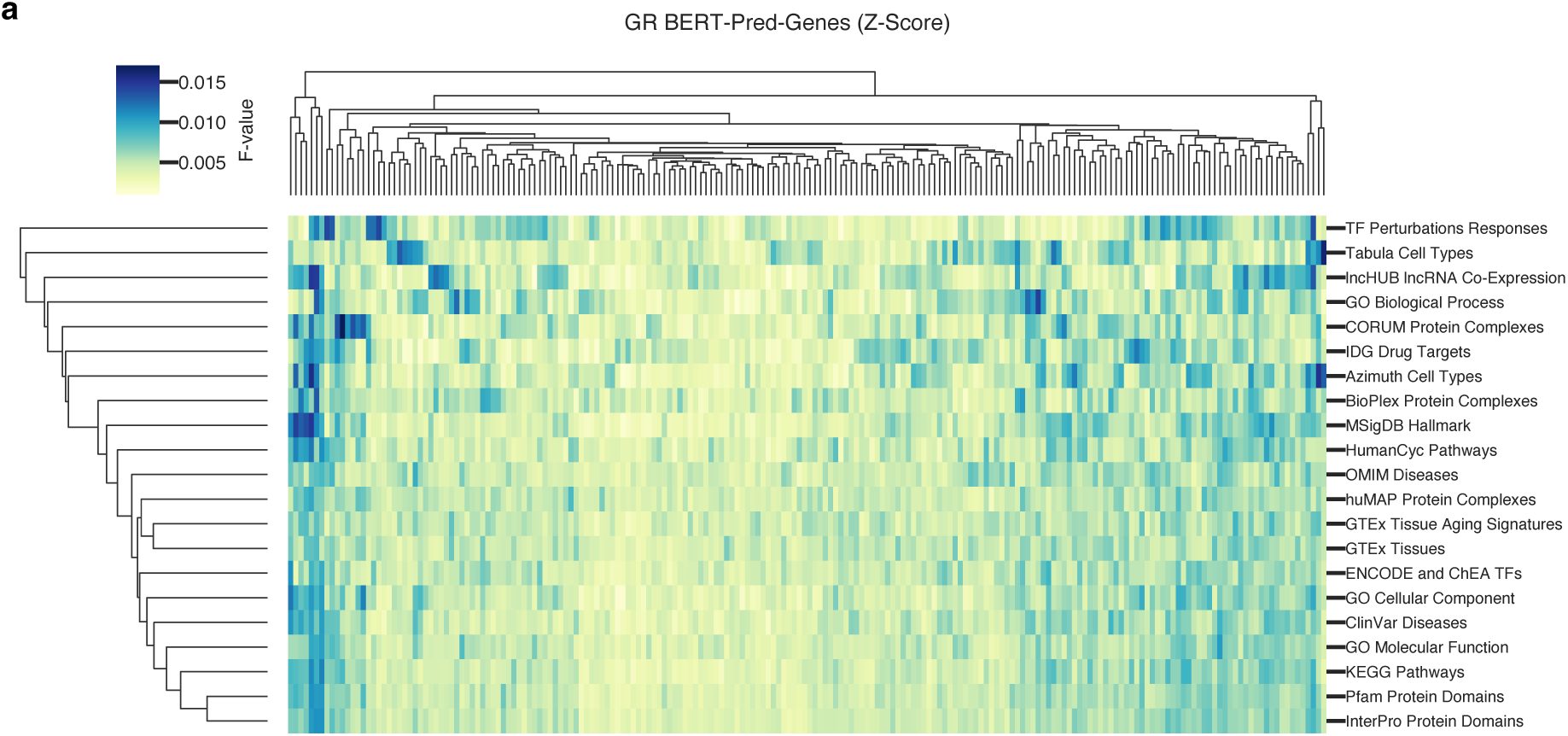
| Dimensional contributions in gene embeddings to gene attribute learning. This figure depicts the relative importance of each embedding dimension in clustering biological attributes of genes, as indicated by the color intensity representing the normalized F-statistic from ANOVA. **a,** Contribution distribution in the BERT-Pred-Genes model using Z-Score-based normalization, with each row corresponding to gene attributes from a specific database and columns denoting embedding dimensions.

**Extended Data Fig. 8.**
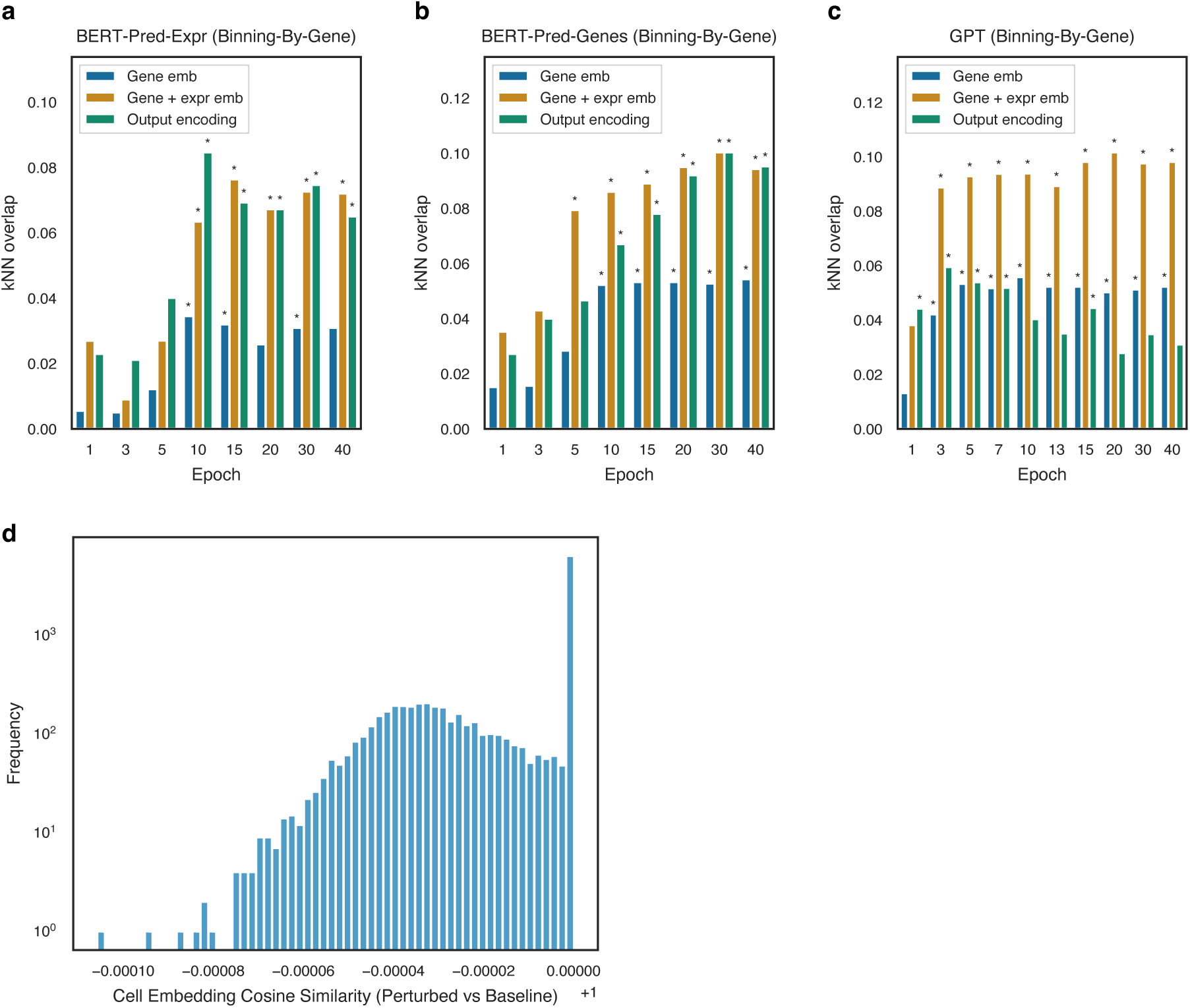
| Model encoding capabilities in reflecting genetic perturbation responses. **a**, Average overlap between the 5 nearest neighbors in the model’s encoding space and 5 genes causing similar transcriptomic alterations was used to measure the model’s ability in reflecting genetic perturbation responses. This figure shows the observed overlap across training epochs in BERT-Pred-Expr model **b,** in the BERT-Pred-Genes model, **c,** and in the GPT model. Results are based on data normalized using the ‘Binning-By-Gene’ method. * *p* < 0.01, as determined by a permutation test with 100 iterations. **d**, Distribution of cosine similarity between cell embeddings from baseline cells and their *in silico* knockdown counterparts (perturbed cells) predicted by the BERT-Pred-Genes model.

**Extended Data Fig. 9.**
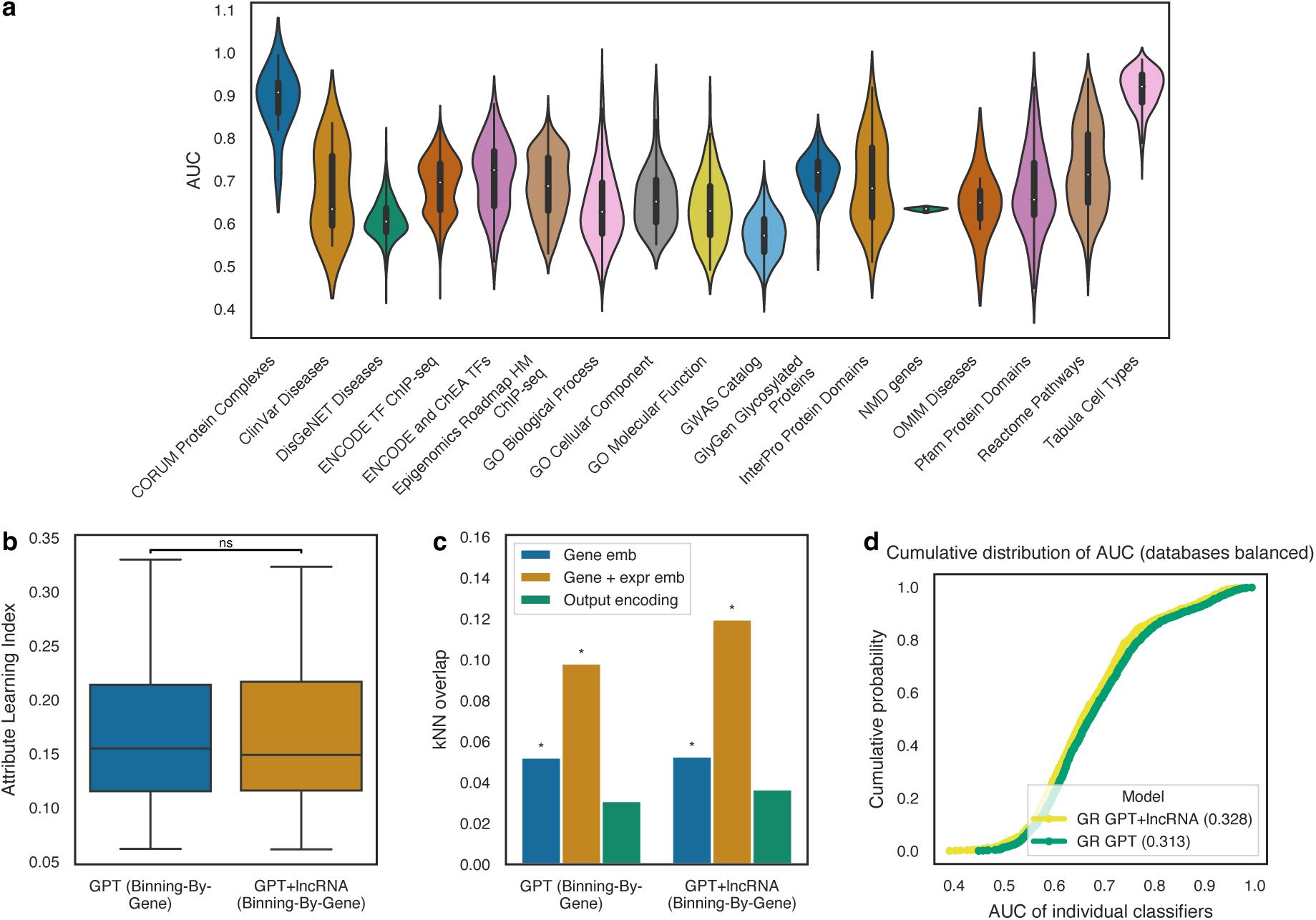
| Classifier performance across gene attributes and in model with lncRNAs. **a**, This figure presents a violin plot illustrating the distribution of AUCs across gene attributes, which are derived from classifiers using embedding in GPT Binning-By-Gene model as features for training. Each classifier was designed to distinguish a target gene class against all other genes. The data visualized here were compiled without balancing the number of classifiers from different databases. **b,** Compares the gene Attribute Learning Index distribution between the GPT (Binning-By-Gene) model incorporating primarily protein-coding genes and the model that includes both protein-coding genes and lncRNA genes. In the boxplot, center line denotes the median; box limits represent the upper and lower quartiles; whiskers extend to 1.5× interquartile range; outliers are depicted as points. ‘ns’, not significant, two-sided paired t-test, after confirming the normality of the data with Shapiro-Wilk test. **c,** Shows the distributions of observed kNN overlaps for both models. * *p* < 0.01, as determined by a permutation test with 100 iterations. **d,** Illustrates the cumulative performance of classifiers using encodings from the two models, with AUC values calculated predominantly using protein-coding genes. *P* = 0.0074, two-sided Kolmogorov-Smirnov test.

**Extended Data Fig. 10.**
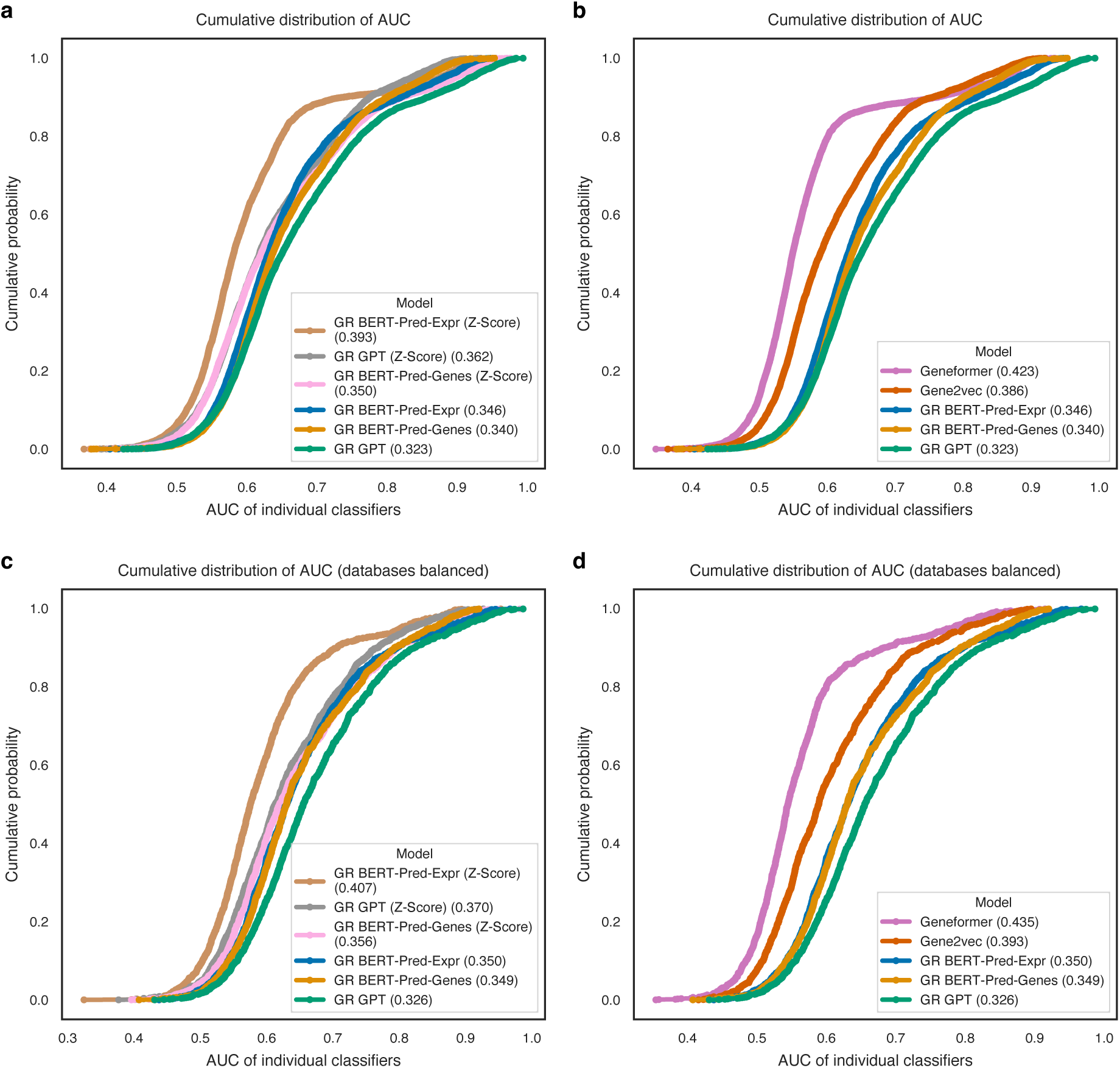
| Cumulative performance of all classifiers using various model encodings. **a** and **b** contrast with Fig. 6 by providing a comprehensive analysis of classifier performance without balancing the number of classifiers from different databases. **c** and **d** contrast with Fig. 6 by classifiers being tasked with distinguishing a target gene class against other classes within the same database (e.g., genes associated with cardiomyopathy vs. genes associated with other diseases) rather than against all other genes (e.g., cardiomyopathy genes vs. non-cardiomyopathy genes). The plots display the cumulative distribution of the AUC values for these classifiers. The x-axis represents the AUC value, while the y-axis indicates the cumulative probability. Points on the plot represent the proportion of classifiers with AUC values less than or equal to the corresponding x-axis value. **a** and **c,** Cumulative distributions of AUCs for classifiers using encodings from models employing different normalization methods. The numbers in parentheses denote the area under the cumulative distribution curve, where lower values signify better classifier performance. In comparisons between Binning-By-Gene and Z-score normalization, all pairs have p-values < 0.0005, two-sided Kolmogorov-Smirnov test. **b** and **d,** Cumulative distributions of AUCs for classifiers using gene embedding from GR models using Binning-By-Gene normalization methods, Geneformer and Gene2vec. For all pairs of comparisons, p-values are < 1E-5, except for GR BERT-Pred-Genes vs. BERT-Pred-Expr in **d**, which has a p-value of 0.50, two-sided Kolmogorov-Smirnov test.

