## Supplementary Information for "Multifaceted Representation of Genes via Deep Learning of Gene Expression Networks"

**Supplementary Table 1. Parameters distribution in the BERT-Pred-Genes model**

| Layer | Num of parameters |
|---|---|
| Predictions | 65,050 (0.9%) |
| Encoder | 2,895,600 (41.7%) |
| Expression embeddings + other embedding params (e.g., layer norm, dropout) | 410,400 (5.9%) |
| Gene embeddings* | 3,577,600 (51.4%) |
| Total | 6,948,650 |

* Gene embeddings for genes not present in the input dataset are excluded from the parameter count calculation.

**Supplementary Table 3. Databases used in gene Attribute Learning Index calculation.**

| Database name | Category | Database ID in Enrichr | Reference PubMed ID |
|---|---|---|---|
| ClinVar Diseases | Diseases/Phenotypes | ClinVar_2019 | 31777943 |
| CORUM Protein Complexes | Protein interactions | CORUM | 36382402 |
| ENCODE and ChEA TFs | Gene expression regulation | ENCODE_and_ChEA_Consensus_TFs_from_ChIP-X | 22955616 20709693 |
| GO Biological Process | Gene ontology | GO_Biological_Process_2023 | 36866529 |
| GO Cellular Component | Gene ontology | GO_Cellular_Component_2023 | 10802651 |
| GO Molecular Function | Gene ontology | GO_Molecular_Function_2023 | 10802651 |
| InterPro Protein Domains | Protein domains | InterPro_Domains_2019 | 36350672 |

| | | | |
|---|---|---|---|
| OMIM Diseases | Diseases/Phenotypes | OMIM_Disease | 25428349 |
| Pfam Protein Domains | Protein domains | Pfam_Domains_2019 | 30398656 |
| Tabula Cell Types | Cell types | Tabula_Sapiens | 35549404 |
| Azimuth Cell Types | Cell types | Azimuth_2023 | 34062119 |
| BioPlex Protein Complexes | Protein interactions | BioPlex_2017 | 28514442 |
| GTEx Tissue Aging Signatures | Aging/Tissues | GTEx_Aging_Signatures_2021 | 32913098 |
| GTEx Tissues | Tissues | GTEx_Tissues_V8_2023 | 32913098 |
| HumanCyc Pathways | Biological pathways | HumanCyc_2016 | 15642094 |
| huMAP Protein Complexes | Protein interactions | huMAP | 33973408 |
| IDG Drug Targets | Drug targets | IDG_Drug_Targets_2022 | 27789690 |
| KEGG Pathways | Biological pathways | KEGG_2021_Human | 27899662 |
| lncHUB lncRNA Co-Expression | lncRNA Coexpression | lncHUB_lncRNA_Co-Expression | 36869839 |
| MSigDB Hallmark | Biological pathways | MSigDB_Hallmark_2020 | 26771021 |
| TF Perturbations Responses | Perturbation responses | TF_Perturbations_Followed_by_Expression | 23193258 |

#### Supplementary Table 5. Databases used for classifier training.

| Database name | Category | Database ID in Enrichr | Min num genes in a class | Num of classes | Num of classes (balanced) | Reference PubMed ID |
|---|---|---|---|---|---|---|
| ClinVar Diseases | Diseases/Phenotypes | ClinVar_2019 | 30 | 11 | 11 | 31777943 |
| CORUM Protein Complexes [#] | Protein interactions | CORUM | 30 | 12 | 12 | 36382402 |
| DisGeNET Diseases | Diseases/Phenotypes | DisGeNET | 100 | 1,072 | 100 | 31680165 |
| ENCODE and ChEA TFs | Gene expression regulation | ENCODE_and_ChEA_Consensus_TFs _from_ChIP-X | 50 | 104 | 100 | 22955616 20709693 |
| ENCODE TF ChIP-seq | Gene expression regulation | ENCODE_TF_ChIP-seq_2015 | 50 | 816 | 100 | 22955616 |

| | | | | | | |
|---|---|---|---|---|---|---|
| Epigenomics Roadmap HM ChIP-seq | Gene expression regulation | Epigenomics_Roadmap_HM_ChIP-seq | 50 | 348 | 100 | 25693563 |
| GlyGen Glycosylated Proteins # | Protein Glycosylation | GlyGen_Glycosylated_Proteins_2022 | 50 | 87 | 87 | 31616925 |
| GO Biological Process | Gene ontology | GO_Biological_Process_2023 | 50 | 739 | 100 | 36866529 |
| GO Cellular Component | Gene ontology | GO_Cellular_Component_2023 | 50 | 131 | 100 | 10802651 |
| GO Molecular Function | Gene ontology | GO_Molecular_Function_2023 | 50 | 137 | 100 | 10802651 |
| GWAS Catalog | Diseases/Phenotypes | GWAS_Catalog_2023 | 50 | 478 | 100 | 36350656 |
| InterPro Protein Domains # | Protein domains | InterPro_Domains_2019 | 30 | 71 | 71 | 36350672 |
| NMD genes | Diseases/Phenotypes | Neuromuscular disease genes (from www.cardiodb.org) | 50 | 2 | 2 | 25589632 |
| OMIM Diseases | Diseases/Phenotypes | OMIM_Disease | 30 | 12 | 12 | 25428349 |
| Pfam Protein Domains # | Protein domains | Pfam_Domains_2019 | 30 | 83 | 83 | 30398656 |
| Reactome Pathways | Biological pathways | Reactome_2022 | 50 | 477 | 100 | 34788843 |
| Tabula Cell Types | Cell types | Tabula_Sapiens | 50 | 469 | 100 | 35549404 |

### Excluded from lncRNA attribute prediction

**Supplementary Table 10. Examples of predicted disease-associated lncRNAs and their supporting literature evidence.**

| Disease | Predicted disease-associated lncRNA | Literature summary of disease-lncRNA association | References |
|---|---|---|---|
| Atherosclerosis | MIAT | MIAT exacerbates atherosclerosis by modulating the miR-181b/STAT3 axis to promote cellular proliferation and inhibit apoptosis, and by sponging miR-149-5p to upregulate CD47, thereby inhibiting efferocytosis and enhancing necrotic core formation in plaques. | 84-86 |
| Colorectal Neoplasms | MIR100HG | MIR100HG stabilizes TCF7L2 mRNA via m6A modification in interaction with hnRNPA2B1, promoting colorectal cancer progression, while β-catenin transcriptionally represses MIR100HG expression through HDAC6-mediated histone deacetylation, affecting cell cycle regulation. | 87,88 |

| | | | |
|---|---|---|---|
| Colorectal Neoplasms | ELFN1-AS1 | ELFN1-AS1 promotes colorectal cancer progression by sponging miR-4644 to upregulate TRIM44, enhancing proliferation and migration, and by recruiting EZH2 and FOXP1 to epigenetically silence TPM1, further driving tumor growth. | 89-92 |
| Colorectal Neoplasms | LUCAT1 | LUCAT1 drives colorectal cancer tumorigenesis by binding UBA52 to activate the RPL40-MDM2-p53 pathway and by antagonizing Nucleolin to enhance MYC expression, thereby promoting cell proliferation and inhibiting apoptosis. | 93,94 |
| Esophageal Neoplasms | HOTAIR | HOTAIR promotes esophageal squamous cell carcinoma progression by enhancing epithelial-mesenchymal transition through upregulating β-catenin and Snail, sponging miR-130a-5p to upregulate ZEB1, and competitively binding miR-204 to modulate expression of HOXC8. | 95-100 |
| Esophageal Neoplasms | HNF1A-AS1 | HNF1A-AS1 promotes esophageal squamous cell carcinoma progression by sponging miR-214 to upregulate SOX-4 and by inhibiting the miR-298/TCF4 axis, thereby enhancing tumor growth, metastasis, and maintaining stemness and epithelial-mesenchymal transition characteristics. | 101-103 |
| Glioma | LINC00470 | LINC00470 promotes glioma cell proliferation and invasion by sponging miR-134 to upregulate MYC and ABCC1, enhances ELFN2 expression through miR-101 sequestration, and modulates autophagy by altering PTEN and AKT signaling pathways. | 104-108 |
| Myocardial Infarction | KCNQ1OT1 | KCNQ1OT1 exacerbates myocardial infarction by sponging miR-466k and miR-466i-5p to upregulate Tead1, enhancing cardiomyocyte apoptosis, and by sequestering miR-26a-5p to regulate ATG12-mediated autophagy. | 109-111 |
| Myocardial Infarction | H19 | H19 ameliorates myocardial infarction by modulating the miR-22-3p/KDM3A pathway to reduce myocardial apoptosis and inflammation, thereby improving cardiac function and remodeling. | 112-115 |
| Non-Small Cell Lung Carcinoma | LINC01234 | LINC01234 promotes non-small cell lung carcinoma progression by sponging various miRNAs to regulate oncogenic targets like OTUB1, VAV3, and GRB2, and interacting with HNRNPA2B1 to modulate miR-106b biogenesis, thereby activating downstream effectors such as c-Myc. | 116-119 |
| Non-Small Cell Lung Carcinoma | LINC01123 | LINC01123 promotes non-small cell lung carcinoma by activating the c-Myc oncogene through the miR-199a-5p/c-Myc axis to enhance cellular proliferation and aerobic | 120,121 |

| | | | |
|---|---|---|---|
| | | glycolysis, and by regulating the miR-4766-5p/PYCR1 axis to drive oncogenic processes in lung adenocarcinoma. | |
| Osteosarcoma | DANCR | DANCR promotes osteosarcoma progression by sponging miR-33a-5p to upregulate AXL and activate the PI3K-Akt signaling pathway, enhancing cancer stem cell features, and by regulating the miR-149/MSI2 and miR-335-5p/ROCK1 axes to facilitate tumor proliferation and metastasis. | 122-124 |
| Prostatic Neoplasms | NORAD | NORAD promotes prostate cancer progression by inducing cell proliferation and migration and facilitating extracellular vesicle release via miR-541-3p-regulated PKM2, contributing to bone metastasis. | 125,126 |
| Stomach Neoplasms | FEZF1-AS1 | FEZF1-AS1 promotes gastric cancer progression by repressing p21 expression through LSD1-mediated H3K4me2 demethylation, activating Wnt signaling, and enhancing cancer stem cell properties via the miR-363-3p/HMGA2 axis. | 127-129 |
| Stomach Neoplasms | MNX1-AS1 | MNX1-AS1 promotes gastric cancer progression by enhancing cell proliferation, migration, and invasion, reducing E-cadherin expression, and increasing levels of PCNA, N-cadherin, vimentin, and MMP-9, potentially through repressing CDKN1A and modulating TEAD4-mediated suppression of BTG2 and activation of BCL2. | 130-132 |
